# Automated analysis of internally programmed grooming behavior in *Drosophila* using a *k*-nearest neighbors classifier

**DOI:** 10.1101/166561

**Authors:** Bing Qiao, Chiyuan Li, Victoria W. Allen, Mimi M. Shirasu-Hiza, Sheyum Syed

## Abstract

Despite being pervasive, the control of programmed grooming is poorly understood. We have addressed this gap in knowledge by developing a high-throughput platform that allows long-term detection of grooming in the fruit fly *Drosophila melanogaster*. Automatic classification of daily behavior shows flies spend 30% of their active time grooming. We show that a large proportion of this behavior is driven by two major internal programs. One of these programs is the circadian clock that modulates rhythms in daily grooming. The second program depends on *cycle* and *clock* and regulates the amount of time flies spend grooming. This emerging dual control model of programmed grooming in which one regulator controls the timing and another controls the duration, resembles the well-established two-process regulatory model of fly sleep. Together, our quantitative approach in *Drosophila* has revealed that grooming is an important internally driven behavior under the control of two regulatory programs.

## Introduction

Grooming is broadly defined as a class of behaviors directed at the external surface of the body. Most animals spend considerable time grooming (Mooring, Blumstein, & Stoner, 2004; Sachs, 1988) and this near universality suggests that grooming likely fulfills an essential role for animals (Spruijt, van Hooff, & Gispen, 1992). Grooming assumes a variety of forms in different species— for instance, birds preen the oily substance produced by the preening gland from their feathers and skin, cats and dogs lick their fur, and flies sweep their body parts with their legs. Though in most cases the primary function of grooming is to maintain a clean body surface, different species-specific forms of grooming have roles in diverse functions such as thermoregulation, communication and social relationships (Ferkin, Leonard, Heath, & Paz-y-Miño, 2001; Geist, Valerius. Walther, 1974; McKenna, 1978; Patenaude & Bovet, 1984; Richard & Dawkins, 1976; G. Schino, 2001; Gabriele Schino, Scucchi, Maestripieri, & Turillazzi, 1988; Seyfarth, 1977; Spruijt et al., 1992; Thiessen, Graham, Perkins, & Marcks, 1977; Walther, 1984).

Though grooming is widely observed and involved in many functions, the basic mechanisms regulating this behavior are still not well understood. Other major behaviors, such as locomotion, are controlled both by external stimuli (stimulated behavior) and by internal programs (programmed behavior). An example of stimulated locomotor activity might be an abrupt evasive response triggered by the sudden appearance of a predator, while programmed locomotor activities, such as daily foraging for food, are essential to maintain vital functions of the organism (Bergman, Schaefer, & Luttich, 2000). Limited data from mammals reveal that grooming, like locomotion, is likely controlled by both external stimuli and internal programs (Hart, Hart, Mooring, & Olubayo, 1992; Hawlena, Bashary, Abramsky, Khokhlova, & Krasnov, 2008; Mooring & Samuel, 1998). However, a detailed understanding of these control mechanisms will require studies in an organism that permits genetic and neural access.

The fruit fly *Drosophila melanogaster* is an ideal model organism with which to dissect the fundamental mechanisms of grooming and its relationship to other behaviors. The fly is known to be a frequent groomer with a rich repertoire of behaviors and a sophisticated genetic toolkit developed to study them (Connolly, 1968; Owald, Lin, & Waddell, 2015). The study of *Drosophila* grooming can be traced back to the 1960’s (Connolly, 1968; Szebenyi, 1969) and notable progress has since been made on the regulation of grooming that ensues immediately after parts of the insect exterior are stimulated with dust particles (Hampel, Franconville, Simpson, & Seeds, 2015; Seeds et al., 2014). While these and most grooming studies thus far have focused on stimulated grooming, understanding mechanisms responsible for programmed grooming will not only identify components distinct to each but also inform us about how programmed grooming is prioritized with regards to other programmed behaviors like locomotion, feeding and sleep.

A major hurdle in detecting programmed grooming in *Drosophila* is the lack of practical methodology. In many cases, fly grooming events are extracted by eye (King et al., 2016; Phillis et al., 1993; Yanagawa, Guigue, & Marion-Poll, 2014). Consequently, these data report only conspicuous behaviors and last for short durations. To improve resolution and accuracy, a number of sophisticated video-tracking methods have been recently developed for fly behavior (Kain et al., 2013; Mendes, Bartos, Akay, Márka, & Mann, 2013). However, these approaches are not ideal for detecting grooming since they focus on leg movements while grooming in flies also entails frequent movements of the antennae, wings and thorax (Seeds et al., 2014). Additionally, the methods are optimized for short-term monitoring (Branson, Robie, Bender, Perona, & Dickinson, 2009; Kabra, Robie, Rivera-Alba, Branson, & Branson, 2013) whereas continuous multi-hour measurements are necessary to dissect fly grooming in relation to other time-dependent behaviors like locomotion and sleep.

To overcome limitations in current methods, we developed a new platform for long-term video-tracking and automated analysis of fly grooming. The layout of our hardware takes advantage of a design widely used in fly locomotion and sleep studies (Gilestro, 2012; Pfeiffenberger, Lear, Keegan, & Allada, 2010) and extends it to studies of grooming in this insect. Our algorithm maps fly activity onto a three-dimensional behavioral space and utilizes *k-*nearest neighbors (*k*NN) method, a machine learning technique, to classify each video frame as grooming, locomotion or rest. Results from multi-day recordings reveal that *Drosophila* spend approximately 30% of awake time grooming and that the temporal pattern of the behavior is tightly regulated by the fly’s internal circadian pacemaker. These findings suggest grooming, similar to feeding and rest, likely serves one or more critical functions in *Drosophila*. Additionally, genetic perturbations and caloric restriction experiments reveal the transcription factors CYCLE and CLOCK as critical parts of an internal program that controls the amount of *Drosophila* grooming. Interestingly, although both *cyc*^01^ and *clk^Jrk^* mutations increase the total amount of basal (internally programmed) grooming, they produce opposite effects when flies are starved (under external stimuli). These grooming data, the easily implementable hardware, and the automated analysis package together permit the construction of high-resolution ethograms of stereotypical fly behavior over the circadian time-scale.

## Results

### Automatic grooming detecting system

To monitor fly behavior, we used a custom-designed system with insects placed individually in tubes with food and cotton at opposite ends (Figure 1A). Tubes were placed in a chamber where temperature and humidity are monitored and controlled. Flies were illuminated from the sides by white light-emitting diodes (LED) to simulate day-night conditions and by infra-red LED from below for video imaging. Videos were captured by a digital camera above the chambers. A sample raw video clip is shown in Video 1.

**Figure 1.**
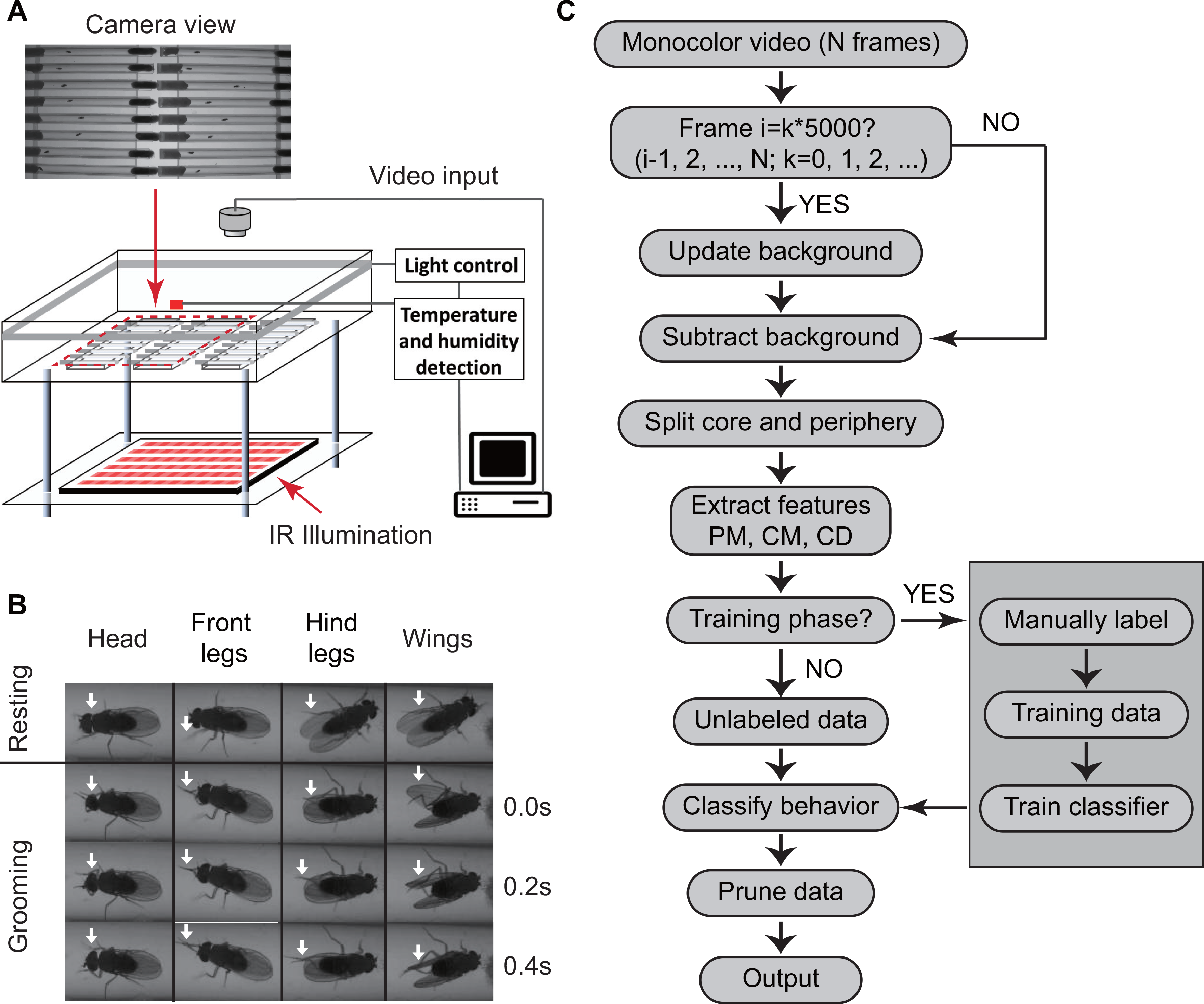
Overview of approach for detecting Drosophila grooming. (A) Apparatus used in recording behavior. Flies constrained to individual tubes, are continuously illuminated by infrared light from below and recorded by a digital camera from above. LED lights on sides of chamber simulate day-night light conditions. Temperature and humidity probes placed in the chamber are monitored by a computer. Inset: A photo of fly tubes in chamber as seen by the camera. (B) Examples of the most commonly observed types of grooming in our experiments. The top row displays postures of a fly in inactive state. The three rows below show how the limbs and body of a fly coordinate to perform specific grooming movements. Arrows point to the moving part during grooming. (C) Flowchart of our algorithm used to classify fly behavior. After generating a suitable background image, the algorithm characterizes movements of fly center (CD), core (CM) and periphery (PM) to fully classify behavior in each frame.

We developed an automated video image analysis package that classifies fly behavior into grooming, locomotion, or rest. Grooming in our algorithm is defined as fly legs rubbing against each other or sweeping over the surface of the body and wings (Szebenyi, 1969) (Video 2, 3), locomotion as translation of the whole body and rest as the lack of either activity. Figure 1B shows images of grooming behaviors frequently observed in our videos involving the head, legs and wings. Since we are primarily interested in detecting grooming events rather than a detailed classification of behavior (Branson et al., 2009), all other behaviors involving body centroid movements are classified as locomotion. This three-tier classification allows our algorithm to efficiently and rapidly interpret grooming events in the recordings without incurring any significant errors in reporting locomotion and rest (see Methods).

To classify behavior, raw videos were processed through four major steps: fly identification, feature extraction, classifier training (optional), and behavior classification (Figure 1C).

### Behavior classification algorithm

Fly identification was accomplished with the following analysis. Flies were first detected in a video frame by computing the difference between the current frame and a reference frame. The reference or background frame was created by comparing two randomly selected frames and erasing all moving objects from one of them (see Methods). We updated the background frame every 1000 seconds to account for changes in the fly’s surroundings (i.e., decrease in the level of food and accumulation of debris within the tube) over the course of multiple hours.

Our algorithm next extracted specific features to classify fly behavior. The features we used are: (1) periphery movement (PM), which characterizes movements of the legs, head and wings; (2) core movement (CM), which quantifies movements of the thorax and abdomen; and (3) centroid displacement (CD), which quantifies whole body displacement. While these intuitive features (PM, CM and CD) are not strictly orthogonal, comparison with orthogonal vectors demonstrated that use of PM, CM, and CD does not compromise accuracy of our algorithm (see Methods and Supplementary Figure S1H). We therefore used PM, CM and CD as our key features throughout the rest of this work. As shown in Figure 2A, relative metrics of PM and CM were different depending on the type of behavior. Specifically, during grooming, the periphery moved more than the core (Figure 2A, top-left, top-right); during locomotion, both parts moved significantly (Figure 2A, bottom-right); while during rest, no significant movement was seen either in the periphery or the core (Figure 2A, bottom-left). The behavior-dependent changes of these features suggest that PM, CM and CD are appropriate metrics for behavior classification. Since differences in fly size can affect values of PM, CM and CD, we also normalized these features to individual fly size before proceeding with further analysis (see Methods).

**Figure 2.**
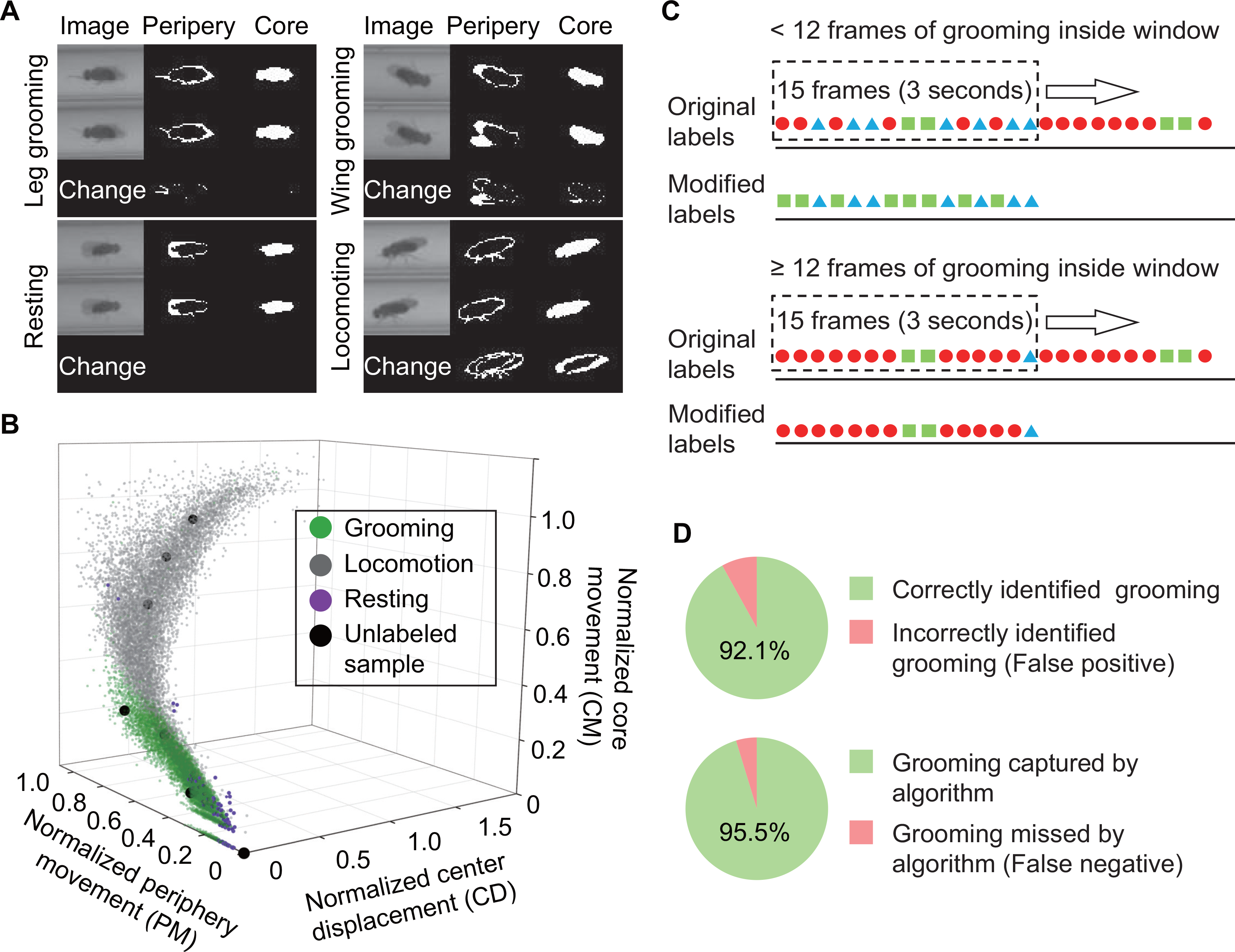
Feature extraction and behavior classification. (A) Examples of original and processed images of a fly displaying different behaviors: Top, left: front leg grooming; top, right: wing grooming; bottom, left: resting; bottom, right: locomoting. In each panel, original images from two consecutive frames are shown on left, periphery in the middle and core on the right. Changes of periphery and core are shown in the bottom row. PM and CM denote differences in the number of pixels representing the fly periphery and core, respectively, in two frames. Features PM and CM are different for different behaviors. Rubbing of front legs manifests through PM (top, left) while sweeping wings affects PM and CM (top, right) (A) Examples of original and processed images of a fly displaying different behaviors: Top, left: front leg grooming; top, right: wing grooming; bottom, left: resting; bottom, right: locomoting. In each panel, original images from two consecutive frames are shown on left, periphery in the middle and core on the right. Changes of periphery and core are shown in the bottom row. PM and CM denote differences in the number of pixels representing the fly periphery and core, respectively, in two frames. Features PM and CM are different for different behaviors. Rubbing of front legs manifests through PM (top, left) while sweeping wings affects PM and CM (top, right) (B) *k-*nearest neighbors (*k*NN) algorithm works by placing an unclassified sample (black circle) representing a frame into a feature space with pre-labeled samples (red/green/blue circles, the training set). The label of the unclassified point is decided by the most frequent label among its *k-*nearest neighbors. The three axes of the feature space are normalized periphery movement (PM), core movement (CM), and center displacement (CD). Fly activity in the feature space is separated into three regions: grooming (red), locomotion (green) and resting (blue). Training samples (N=18000 for each color) and 9 unlabeled samples in PM-CM-CD space are shown. (C) Grooming data are pruned after identification by the *k*NN classifier. A frame is finally labeled as grooming only if this frame is in a group of 15 frames in which 12 or more were labeled as grooming by the classifier. Frame previously labeled as grooming by the classifier but that did not pass the pruning procedure is relabeled as locomotion. (D) (Top) 92% time of all grooming detected by the program is correct. We randomly sampled 10% of all grooming classified by our algorithm in an eight-hour video, and then manually determined the false positive rate by watching the video. The false detection (red) results from movements that are similar to grooming, such as slow body displacement and bending of the abdomen and mouth. (Bottom) Our algorithm successfully detects 95.5% of all grooming in a video (bottom). The circle represents all the grooming in a 460 minutes video and the green area represents grooming detected by the program.

We then classified fly behavior by applying the *k-*nearest neighbors (*k*NN) technique to the normalized features (Bishop, 2007). Briefly, *k*NN works by placing an unlabeled sample into a feature space with pre-labeled samples serving as a training set for the algorithm. The label or class of the unlabeled sample is then decided by the label that is most common among its *k-*nearest training samples. In our case, the nearest neighbors were searched through a *k-*d tree algorithm (Sproull, 1991). To construct the *k*NN classifier, we prepared a training set by visually labeling fly behavior from 25000 frames and mapping them onto a three-dimensional feature space where the axes correspond to PM, CM and CD (Figure 2B, color symbols). We tested values of the parameter *k* between 1 and 50 and settled on *k*=10 to achieve balance between computing time and accuracy (see Methods).

Finally, we pruned output labels from the *k*NN classifier (Figure 2C). The algorithm calculates features from every two consecutive frames, resulting in some classifications being confounded by short-term fly activity. For example, features extracted from only two frames often cannot distinguish a fly stretching its body parts from one that is grooming. Based on our observations during creation of the training set, a typical grooming bout lasts >3 seconds or for 15 frames at our normal frame rate, longer than an average stretching event, which lasts for ~1 second. Accordingly, we applied a 15-frame-long temporal filter that slides one frame at a time to eliminate false grooming labels caused by short, grooming-like behavior. Grooming designations were retained only if at least 12 grooming frames are found within the window. Otherwise, all grooming frames were relabeled as locomotion once the left edge of the window reaches the fifteenth frame (Figure 2C). These pruned labels were the final output of our grooming classification algorithm.

The accuracy of our algorithm was evaluated by comparing the computer-identified grooming with manually-labeled grooming identified by visual inspection. We tested a total of 8 hours of videos, including 15 individual flies (see Methods), and found that of the grooming events picked out by our algorithm, 92.1% were manually verified as true grooming events (Figure 2D, top panel). Furthermore, among all manually scored grooming events, 95.5% were successfully identified by our computational method (Figure 2D, bottom panel). These test results suggest that our method identifies grooming with high fidelity.

### Grooming plays an important role in the daily life of Drosophila

To determine how grooming is coordinated within the 24-hr period, we examined fly behavior over the course of several days in 12 hour light: 12 hour dark (LD) conditions (Figure 3). In LD cycles (for constant darkness, see Figure S2A), locomotion levels showed the familiar morning (M) and evening (E) peaks around the time lights turn on and off (Figure 3A middle), respectively (Schlichting et al., 2016; Stoleru, Peng, Agosto, & Rosbash, 2004). Nearly coincident with increases in locomotion were increases in fly grooming (Figure 3A bottom), although these time-dependent peaks in grooming were more subdued compared to those in locomotion. While basal locomotion during mid-day or night decreases to < 5% of the M/E peak values, basal grooming during the same duration was maintained at ~14% of the peak values (Figure 3A, rectangles). The smaller time-dependent variations in grooming resulted from 20-40 bouts per hour with the longest pause between two bouts being ~83 minutes on average (Figure 3B). In contrast, the longest pause between two consecutive bouts of locomotion was ~116 minutes (Figure 3B). Because grooming bouts were on average shorter than locomotion (Figure 3C), a typical fly under LD conditions spent approximately 9% of its daily time grooming, compared to 20% of time in locomotion (Figure 3D). That is, the average fly spends ~30% of its active time grooming. The frequency of grooming behavior suggests that maintenance of a low but steady rate of grooming is important for the animal.

**Figure 3.**
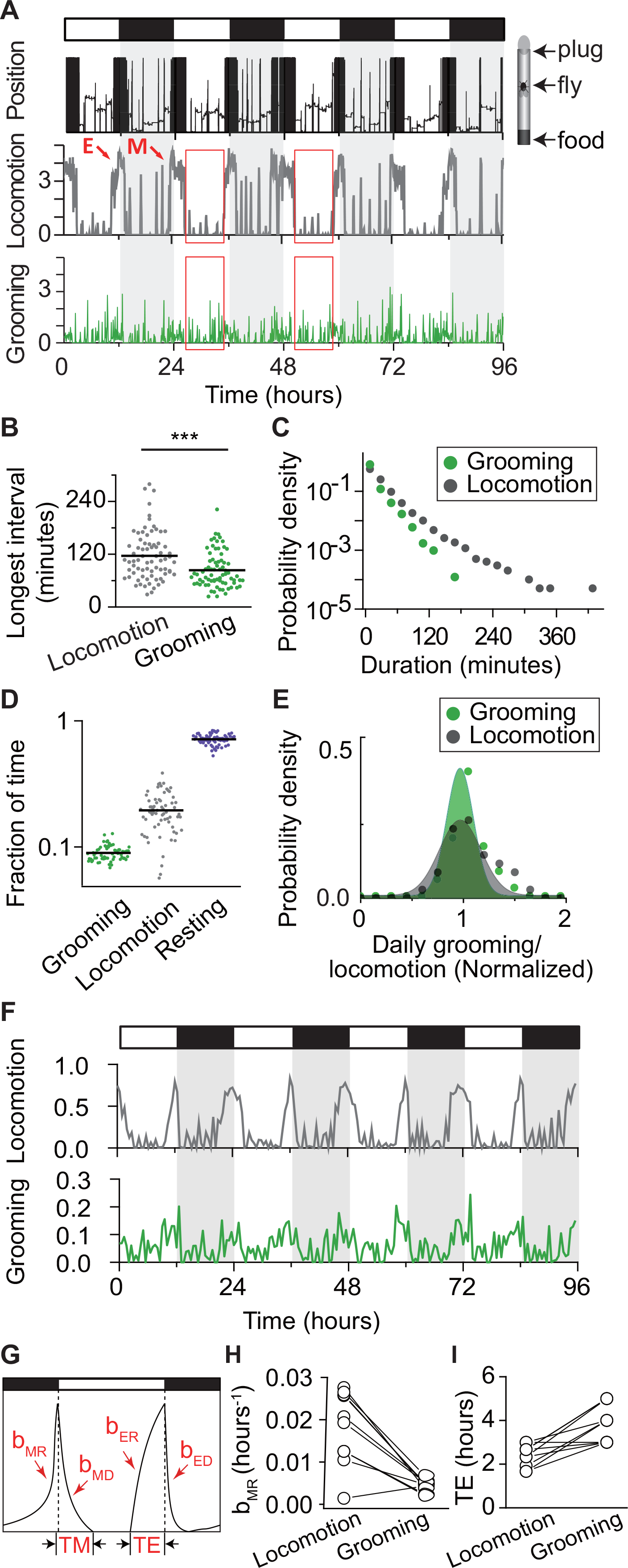
How grooming fits into the daily routine of a fly. (A)Position within the tube (top row), locomotion (middle) and grooming (bottom) of a single fly during four days in LD cycles. Locomotion is shown in terms of the duration (minutes) spent in locomotion in five minute bins. Morning and evening peaks in locomotor activity are marked as M and E. Grooming is shown in terms of time spent grooming (minutes) in five minute bins. White/black bars indicate light/dark environmental conditions, respectively. (B) Longest intervals between grooming events (green) and between locomotion events (black). Each point represents an individual fly recorded for a day. N= 74 flies, p=7.09×10^-5^ (C) Probability density of the duration of grooming events (green) and locomotion events (black). N= 20 flies. (D) Fraction of time spent in grooming, locomotion and resting states in WT flies. On average, flies spend about 9% of time grooming every day and 20% time in locomotion. N=66 flies. (E) Inter-individual differences in daily grooming and locomotion. Normalized distributions of individual grooming and locomotion (total individual daily grooming or locomotion divided by population average) are fitted to normal distribution functions. Variation in daily grooming time among individuals is significantly less than the variation in locomotion. Standard deviation of grooming is 0.14 compared with 0.34 for locomotion. N=66 flies. (F) Fraction of time spent locomoting and grooming by an individual fly. Fraction is calculated every 30 minutes. (G) Sketch of the mathematical model that uses four normalized exponential terms to describe temporal patterns of a fly activity. Parameters *b*_*MD*_, *b*_*ER*_, *b*_*ED*_, *b*_*MR*_, TM and TE (see text) are marked in the plot. (H), (I) Comparison of *b*_*MR*_ and TE values between locomotion and grooming. Each circle represents an individual fly and data from the same fly are connected by a solid line.

The reduced temporal modulations in individual grooming behavior was accompanied by similarly reduced variability in grooming levels between individual flies (Figure 3E). To compare variability of grooming and locomotion across the population, we constructed normalized distributions for the two behaviors by calculating daily grooming and locomotion times of individuals and dividing these by the respective population means. These data revealed that, under LD conditions, the standard deviations in grooming and locomotion were 0.14 and 0.34, respectively. Similarly, in constant darkness, they were 0.16 and 0.25 (Figure S2B). The relatively low individual variation in grooming behavior suggests a consistent, internally programmed drive to groom. Together, the considerable time spent and the low population-wide variability in grooming are consistent with an important role for this behavior in the daily routine of *Drosophila melanogaster*.

To quantitatively compare the temporal patterns of grooming and locomotion (Figure 3F), we applied a previously developed mathematical function that models fly activity in terms of exponential functions (A. Lazopulo & Syed, 2016). The functions are defined by four rate parameters *b*_*MR*_, *b*_*MD*_, *b*_*ER*_ and *b*_*ED*_, where subscripts denote morning rise (MR), morning decay (MD), evening rise (ER) and evening decay (ED), and two duration parameters that describe the relative durations of morning (TM) and evening (TE) peaks (Figure 3G). We previously proposed that these parameters may reflect kinetics of biochemical substrates underlying the specific fly behavior described by the model (A. Lazopulo & Syed, 2016). We fitted this model to grooming and locomotion of individual wild-type flies for 3-4 days in LD conditions. Results showed that the rate parameter *b*_*MR*_ of grooming was smaller than that of locomotion (8 out of 9 flies, Figure 3H), indicating a slower increase in night-time grooming activity and consistent with a smaller change in grooming between day and night (Figure S3A). Additionally, the evening duration parameter (TE) for grooming was greater than that for locomotion (Figure 3I), indicating that the evening peak in grooming lasted longer. In contrast, the other model parameters did not show significant differences between locomotion and grooming (Figure S3B-E), raising the possibility that, in addition to their differences, the two behaviors may also share some common underlying regulatory substrates.

### Temporal pattern of grooming is under control of the circadian clock

The circadian clock modulates a wide range of fly behaviors (Allada & Chung, 2010), including locomotor activity. To test whether basal grooming is also under circadian control, we monitored grooming in wild-type (WT) and circadian mutants *per^S^, per ^L^*, and *per^0^* for 4 days in LD followed by 4 days in constant darkness (DD, Figure 4A). Mutations of the endogenous circadian clock cause altered circadian period length or arrhythmia in the absence of light stimulation (DD). *per^S^* and *per ^L^* mutants have short and long circadian periods, respectively while *per^0^* mutants are arrhythmic. Population-averaged LD data showed that light was a strong zeitgeber of grooming even for circadian mutants, while the DD data revealed that grooming is circadian-regulated, as circadian mutants exhibited grooming behavior with the expected changes in periodicity (Figure 4A, top three panels) or arrhythmia (Figure 4A, bottom panel). Autocorrelation analysis of wild-type LD data over a few hours showed weaker correlation in grooming compared to locomotor activity (Figure 4B), while spectral analyses showed oscillation periods in constant darkness to be 23.73 ± 1.10 hours, 18.70 ± 0.71 hours, and 28.48 ± 1.13 hours for WT, *per^S^*, and *per ^L^* flies, respectively (Figure 4C). In *per^0^* flies, grooming activity does not show any significant periodicities in spectral analysis (data not shown). These long time-scale oscillatory periods are in agreement with those of locomotor rhythms (Figure S2C, D). The observed shifts in the period of grooming rhythms, consistent with well-characterized molecular perturbations of the clock, suggest that the circadian clock temporally modulates grooming in *Drosophila*. Interestingly, *per^S^, per ^L^*, and *per^0^* mutations cause major changes in temporal grooming rhythms while causing no significant change in the total level of grooming (Figure 4D). This result is consistent with at least two sets of regulatory mechanisms for basal or internally-programmed grooming: circadian regulation to regulate the timing of grooming, and an internal drive to regulate the amount of grooming.

**Figure 4.**
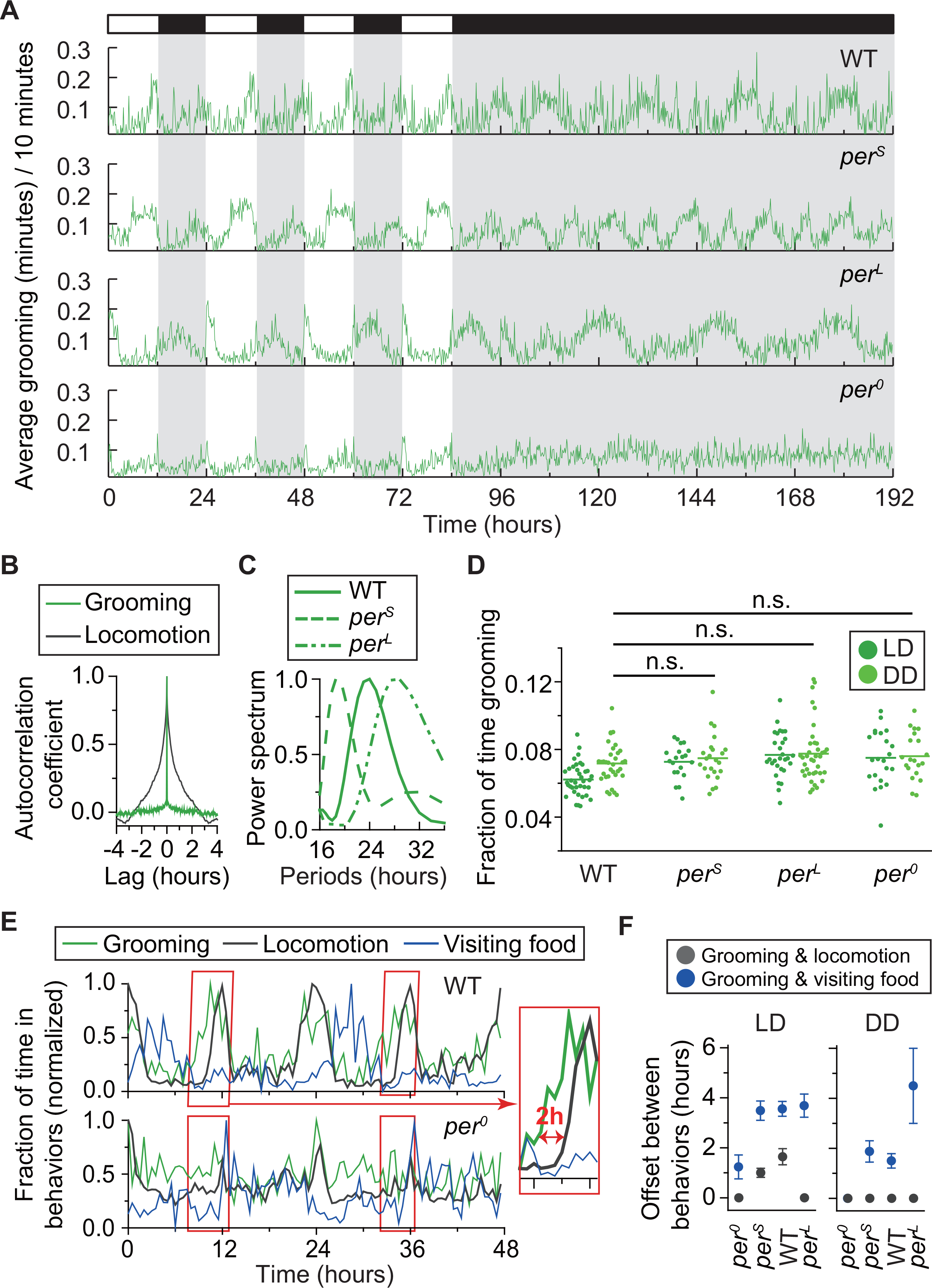
Grooming is under control of the circadian clock. (A) Grooming activity (in 10 minute bins) of wild-type and clock mutants during four days in LD cycle followed by four days in DD cycle. Grooming traces are population averages. In DD, WT grooming continues to show 24 hr rhythms. In comparison, grooming in *per* ^*S*^or *per*^*L*^ flies show shorter or longer rhythms, respectively. For *per*^0^ flies, grooming is arrhythmic in DD. N=8 WT, 8 *per*^*S*^, 10 *per*^*L*^, 10 *per*^0^. (B)Autocorrelation of grooming and locomotion. The relatively rapid drop in correlation among individual grooming events suggest greater short-term (for time lags > 2 minutes) independence of these events when compared to locomotion. N=8 WT flies. (C) Long-term correlation and circadian rhythmicity in grooming shown by average power spectra of wild-type, *per*^*S*^ and *per*^*L*^ flies. N=34 WT, 23 *per*^*S*^, 38 *per*^*L*^. (D) Daily time spent in grooming is generally unaffected by aberrant circadian rhythms. N=34 WT, 23 *per*^*S*^, 38 *per*^*L*^, 20 *per*^0^. In DD, p=0.36 for WT vs *per*^*S*^, p=0.1 for WT vs *per*^*L*^, p=0.23 for WT vs *per*^0^. (E) Normalized average amount time spent in grooming (green), visiting food (blue) and locomotion (gray) during two days in LD (see Methods). Each behavior time series is normalized by its maximum to allow for easy comparison of their relative phases. In wild-type flies (top panel), burst in visiting food happens 2-4 after the morning peak in locomotion. Onset of evening peaks in grooming usually occurs earlier than the peak in locomotion (red boxes). A close up view is shown on right. N = 8 WT flies (top panel) and N = 10 *per*^0^ flies (bottom panel). (F) The time difference in onset of bursts in grooming and locomotion (gray), grooming and visiting food (blue), in LD (left) and DD (right).

Because *Drosophila* feeding activity is also regulated by the circadian clock (Chatterjee, Tanoue, Houl, & Hardin, 2010; Xu, Zheng, & Sehgal, 2008), we tested whether the observed rhythms in grooming could be an indirect effect of rhythmic food intake, with food debris serving as the external stimulus (Hampel et al., 2015; Seeds et al., 2014). Since our assay is not optimized to directly measure feeding, we used prolonged proximity (> 3 seconds, < body length) with food as an indication of feeding behavior (see Methods). This analysis demonstrated that, in LD conditions, wild-type controls exhibited robust oscillations in visits to food with a peak around 3 hours after lights turn on (Figure 4E, top panel, blue). The peak time in contacting food was offset by 2-4 hours from nearby peaks in grooming (Figure 4E, top panel, green). This temporal offset suggests that periodic contact with food is unlikely to be the external stimulus that drives rhythms in basal grooming. Locomotor rhythms are also unlikely to be the primary driver of grooming rhythms since the onset of evening peak in grooming was ~ 2 hours earlier than the evening peak in locomotion (Figure 4E, top panel, red boxes and inset). This is consistent with the comparison in Figure 3I, which shows that the grooming evening peak lasts longer than the locomotion evening peak. These temporal offsets in grooming, feeding and locomotion were typically reduced in constant darkness (Figure 4F) and nearly absent in *per^0^* mutants (Figure 4E, bottom panel; Figure 4F), suggesting that they result from a combined effect of the external zeitgeber and the internal pacemaker. Together, these results suggest that the circadian clock directly influences temporal patterns in grooming, thus identifying endogenous timekeeping as a likely internal program that influences the *Drosophila* grooming circuitry.

### Grooming duration is controlled by cycle and clock

The circadian clock appears to affect mainly the temporal pattern of grooming without altering the total time flies spend in the behavior (Figure 4D). Based on grooming data from other animals implicating the behavior in stress relief (Chen et al., 2010; Hart, 1988; Gabriele Schino et al., 1988), we hypothesized that flies with altered stress response may also exhibit altered levels of daily grooming when exposed to a common external stimulus.

The fly transcription factors CYCLE (CYC) and CLOCK (CLK) activate essential clock genes by binding E box sequences as a heterodimer (Crane & Young, 2014). Although they are best known for maintaining circadian rhythmicity, *cycle* and *clock* have also been implicated in regulating sleep need in response to sleep deprivation and adjusting locomotor output in response to nutrient unavailability (Hendricks et al., 2003; Keene et al., 2010; Shaw, Tononi, Greenspan, & Robinson, 2002). To test if *cycle* or *clock* play a role in setting the level of grooming under normal LD conditions, we measured the behavior in *cyc^01^* (Rutila et al., 1998) and *clk^Jrk^* (Allada, White, So, Hall, & Rosbash, 1998) mutants. The data showed increased daily average grooming in both mutants relative to genetic controls (Figure 5A, B). The shared increase in grooming duration in these flies is accompanied, however, by opposing changes in their locomotion. Relative to their controls, *cyc^01^* flies spent less time, while *clk^Jrk^* flies spent almost twice as much time in locomotion (Figure S4A, B). These results reveal a differential reprioritization of behavioral outputs by the two mutations, similar to phenotypic differences reported previously in sleep studies involving *cyc^01^* and *clk^Jrk^* (Hendricks et al., 2003; Shaw et al., 2002).

**Figure 5.**
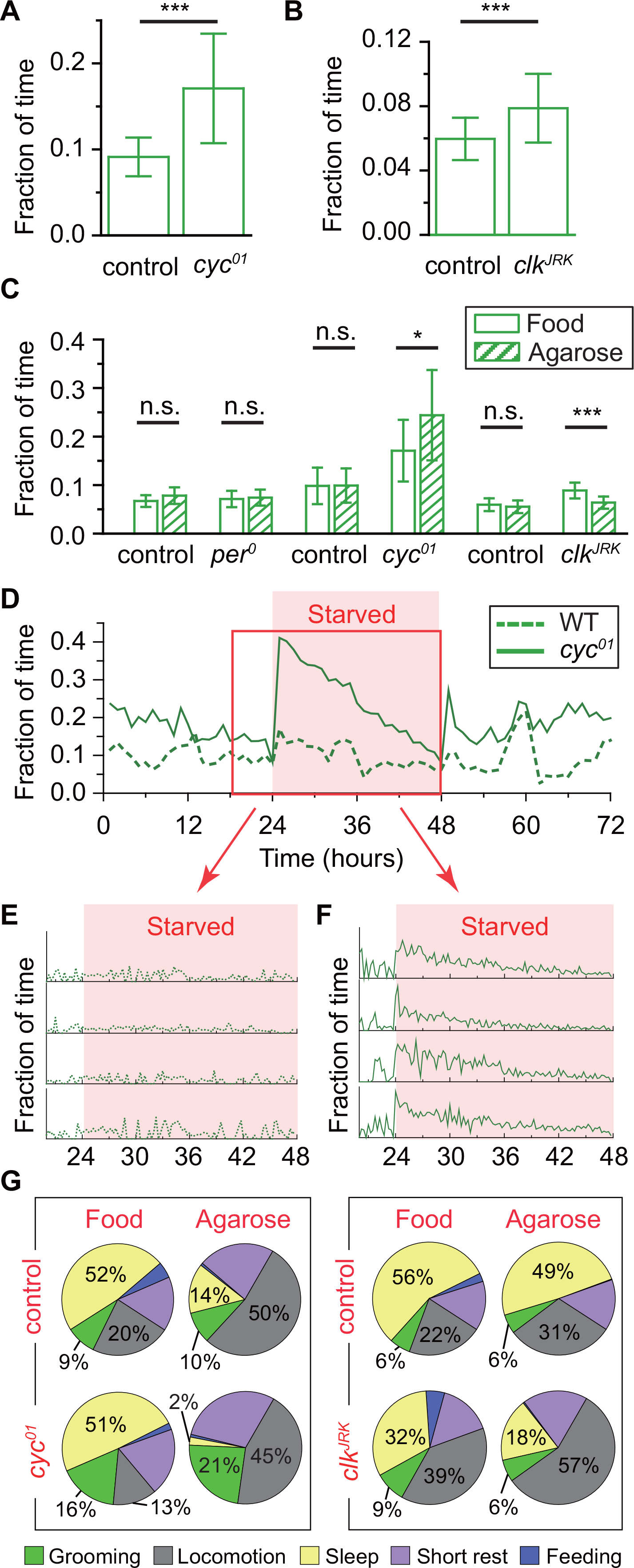
Amount of grooming is controlled by CYCLE and CLOCK. (A) *cyc*^01^ flies groom ~60% more than their background control (p=0.0011). The increase is unlikely to be a result of a non-working clock, as arrhythmic *per* ^0^ flies do not show a similar change (Figure 4C). Instead, lack of CYCLE or genes it helps transcribe, likely elevates baseline grooming. (B) Grooming of *clk^Jrk^* flies and their background control. *clk^Jrk^* flies show significantly more grooming than control (p=7.91×10^-9^). (C) Differential grooming response to stress through starvation. Data are averaged from the second day of 3-day experiments in which during the second day flies were either kept in normal diet (“Food”) or placed in 1% agarose diet (“Agarose”). *cyc*^01^ flies show increased amount of grooming when starved, while *clk^Jrk^* flies groom less during starvation. Other tested genotypes maintain grooming at their respective normal levels. N=18, p=0.567 for *per*^0^ flies and N=20, p=0.09 for control. N=18, p=0.029 for *cyc*^01^ flies and N=14, p=0.554 for control. N=28, p=1.75×10^-6^ for *clk^Jrk^* flies and N=28, p=0.09 for control. (D) Temporal patterns in WT and *cyc*^01^ grooming during a 3-day 12:12 LD experiment in which flies are starved on a 1% agarose diet during the second day (shaded). Population average data plotted in one-hour bins (N=10 WT; N=10, *cyc*^0^). (E) and (F) Examples of individual (E) WT and (F) *cyc*^01^ flies. Individual time-series are binned in 15 minutes and include four hours before the start of starvation. (G) Average fraction of time flies spend in grooming (green), locomotion (gray), sleep (yellow), short rest (purple), and feeding (blue). N=18 *cyc^01^* flies and 14 of control. N=26 *clk^Jrk^* flies and 28 of control.

Because many different types of stress disrupt circadian regulation of locomotor activity, we next tested whether stress also disrupts circadian regulation of grooming behavior. We subjected *per^0^*, *cyc^01^*, *clk^Jrk^* mutants, and their controls to a common stressor: unavailability of nutrients. Previous work had shown that starvation causes loss of circadian regulation (Keene et al., 2010). Flies were allowed to acclimate to standard food and LD cycle for one day, after which grooming was recorded for the next three days with the second day either in normal food or 1% agarose. Consistent with the hypothesis that grooming behavior is circadian-regulated, we found that starvation disrupted circadian oscillations in grooming behavior, as well as locomotor activity, in wild-type flies (Figure S4C, D). Moreover, the starvation-induced disruption of circadian regulation is thought to result from the reprioritization of behavior: flies upregulate locomotor activity and downregulate sleep to engage in starvation-induced foraging behavior that overrides and is independent of circadian regulation (Keene et al., 2010). Consistent with this, all mutants and controls exhibited increased locomotor activity under starvation conditions (Figure S4C).

To test whether this reprioritization of behavior extended to grooming, we examined total levels of grooming under starvation conditions, as measured by total time spent grooming. We expected that grooming behavior would either be deprioritized relative to locomotor activity and down-regulated, similar to sleep, or increased relative to normal nutrient conditions, similar to locomotor activity, because flies are sleeping less and spending more time being active. Unexpectedly, we found that starvation induced no significant change in time spent grooming in both *per^0^* mutants and control animals. This result supports the hypothesis that the daily time spent grooming is regulated by an internal program independent of circadian regulation and suggests that this internal program is resistant to starvation-induced stress.

This reprioritization of behavior is even more dramatic in two other circadian mutants *cyc^01^* and *clk^Jrk^*, both lacking a functional clock. Relative to controls or *per^0^* mutants, both *cyc^01^* and *clk^Jrk^* were previously shown to dramatically downregulate total sleep amount under starvation conditions, presumably by upregulating locomotor activity because of increased metabolic stress (Keene et al., 2010). Consistent with this, we found that *cyc^01^* and *clk^Jrk^* exhibited increased locomotor activity under starvation conditions (Figure S4C). We then tested whether this increase in metabolic stress was sufficient to deprioritize grooming behavior under starvation conditions. In support of this hypothesis, *clk^Jrk^* mutants under starvation conditions exhibited a modest decrease in time spent grooming relative to normal nutrient conditions (Figure 5C). Unexpectedly, however, *cyc^01^* exhibited the opposite response: a significant and robust increase in time spent grooming under starvation conditions. This increase in *cyc^01^* grooming mainly occurs during the first ~10 hours of their introduction to the agarose-diet (Figure 5D-F). There is at least another previously reported case in which *cyc^01^* mutants have a distinct phenotype relative to other circadian mutants: a disproportionately strong rebound in sleep after sleep deprivation, thought to result from defects in heat-shock stress response (Shaw et al., 2002). This suggests that the immediate, excessive grooming in response to starvation as exhibited by *cyc^01^* may also be due to defects in heat-shock stress response in the mutant. Taken together, our data show that while the internal drive to groom is not normally impacted by metabolic stress, the loss of the two circadian clock components *cyc* and *clk* increases the internal drive to groom (Figure 5A,B) and alters the grooming response to starvation conditions (Figure 5C). The opposite responses to starvation by *cyc^01^* and *clk^Jrk^* flies may be due to CLOCK or CYCLE interacting exclusively with partners outside of those they bind as a heterodimer (Hendricks et al., 2003), one consequence of which may be aberrant expression of heat-shock genes in *cyc^01^* but not *clk^Jrk^* flies (Shaw et al., 2002).

To determine to what extent observed changes in grooming and locomotion affected the other behavioral classes, we next broadened our analysis to include rest, feeding, and sleep. Feeding was calculated in terms of extended period spent near food (as defined for Figure 4E) and sleep was determined in terms of prolonged rest, ≥ 5 min episodes of no grooming or locomotion (Shaw et al., 2002). The analysis revealed a general trend across all tested strains: lack of nutrients diminished time spent feeding and sleeping but increased time dedicated to short rests and locomotor activity (Figure 5G and Figure S5). Increase in rest time is surprising since re-allocation of time away from sleep (prolonged rest) time would predict a similar reduction in short rests. That flies instead spend more time resting during starvation implicates a sophisticated energy-balance mechanism that couples increase in locomotor activity, needed for foraging, with increase in short rests, presumably needed to improve efficiency in foraging expeditions.

Despite substantial reduction in sleep under starvation conditions, grooming levels were held approximately constant in all control and *per^0^* flies (Figure S5). This result shows that grooming behavior is prioritized above sleep during starvation, as time spent grooming could otherwise be spent sleeping or foraging. Stability in time spent grooming in the absence of food further supports the contention that much of the grooming detected in our experiments is not stimulated externally by food contact but rather controlled by internal programs. As noted above, lesions in *cyc* and *clk* affected this stability and resulted in elevated grooming (Figure 5A, B). Through the ethograms we found that in case of *cyc^01^*, the increase in grooming came from loss of locomotor activity while in case of *clk^Jrk^* the increase came from loss of sleep (Figure 5G). This result supports the hypothesis, now with more detail, that the *cyc^01^* and *clk^Jrk^* mutations alter the insect’s internal homeostasis in distinct ways, which also helps explain why they exhibit starkly different responses when placed under metabolic stress (Figure 5C, G).

Accumulated data from our experiments suggest that grooming is an innate fly behavior controlled by two major regulators. One of these regulators controls temporal patterns in grooming and another controls amount of time spent in grooming. Circadian genes *per*, *cyc* and *clk* are involved in controlling the timing of peaks/troughs in grooming rhythms while *cyc* and *clk* are also involved in setting how much time is spent grooming. The apparent absence of *per* from the second regulatory mechanism is consistent with the idea that the two control mechanisms are able to operate independently.

## Materials and methods

### Fly strains

Clock mutants *per^S^, per ^L^*, and *per^0^* were backcrossed for five-six generations to an *iso31* with *mini-white* insertion strain. *cyc* mutants, gifts from William Ja (The Scripps Research Institute), have the *Canton S* background. *Clk^Jrk^* flies were backcrossed for five generations to *iso31*. Flies were bred and raised at 23°C and 40% relative humidity on standard cornmeal and molasses food. All experiments were done with 5-8 days old males at 26^0^C and 70-80% relative humidity in a custom-built behavior tracking chamber (Figure 1). For each experiment, control strain refers to the genetic background of a mutant. WT flies in Figure 3 refer to the *Canton S* line.

### Behavior tracking apparatus

#### Chamber

Flies were placed individually in glass tubes (Trikinetics Inc., Waltham, MA, PGT5x65) with food and a cotton plug at opposite ends. Twenty tubes were placed on a custom-designed plate inside a transparent acrylic cuboid box for simultaneous imaging. Temperature and humidity were monitored every 5 mins with a digital thermometer (Dallas Semiconductor, Dallas, TX, DS18B20) and a humidity sensor (Honeywell, Morris Plains, NJ, HIH-4010), respectively, while a wet sponge inside the chamber kept the relative humidity around 70%-80% (Figure S1A).

#### Illumination

The chamber was illuminated by two sets of light-emitting diode (LED) strips. White LEDs (LEDwholesalers, Hayward, CA, 2026) producing ~700 lux were used to simulate daytime conditions and infrared LEDs (LEDLIGHTSWORLD, Bellevue, WA, SMD5050-300-IR 850nm) were used to visualize the flies at all time.

#### Camera

A CCD monochrome camera (The Imaging Source, Charlotte, NC, DMK-23U445) fitted with a varifocal lens (Computar, Cary, NC, T2Z-3514-CS) was used for video imaging. To minimize influence of chamber’s light/dark conditions on video quality, we put a 780 nm long pass filter (Midopt, Palatine, IL, LP780-30.5) in front of the lens. Videos were saved as 8-bit images in .avi format with 1280 x 960 resolution at 10 Hz and down-sampled as needed.

### Analytic hardware and runtime

Using a desktop computer with Intel Core i7-4770 3.4 GHz processer and 4 × 4 G DDR3 1600 MHz RAM, it takes ~7 hours to extract grooming, locomotion and rest data from an 8-hour video of 20 flies recorded in 10 Hz (in total 288000 frames) at 1280 pixel × 960 pixel resolution. Videos are analyzed every 2 frames (5 Hz), which is sufficient to capture grooming events.

### Starvation media

Media for starvation experiments was made by dissolving 1% agarose in water.

### Algorithm for automatic detection of grooming

All computational analyses were done with custom-written Matlab scripts that will be available at http://syedlabmiami.weebly.com/software.html

#### Fly shape extraction

Fly shape was extracted by applying a background subtraction algorithm as described below.

#### Creating Background

The background or reference frame is constructed by randomly picking two frames, a template and a contrast, and comparing their pixel grayscale values and erasing all moving objects from the template frame. To remove the fly from the template frame, we replace the pixels belonging to the fly with corresponding pixels from the contrast frame, relying on the fact that a fly is always darker than the surrounding objects. The template frame with no fly present then becomes the background frame. Additionally, because a fly’s surroundings, including food debris, change substantially during the course of an experiment (Figure S1B), the background frame is regenerated every 1000 seconds. Lastly, if a fly occupies the same area in the template and contrast frames, the overlapping region cannot be erased on the template. To circumvent this problem, every time a background frame is generated, we randomly choose 7, instead of 1, frames as contrast frames and compare all of them with the template. When a fly does not move for more than 1000 seconds, the fly will not be removed from the background and cannot be detected in other frames during this 1000 seconds. Thus when a fly is not detected, we consider the fly to be stationary at the position where it was last detected.

To reduce effects of charge coupled device (CCD) image noise and fluctuations in the system, we set a minimum change *C*_0_ as the threshold to accept grayscale changes from fly movements. We denote the grayscale value of a pixel located at *(x, y)* (in units of pixel, in our case, *x* ∊ [1:1280], *y* ∊ [1:960]) in the template as *I*_*template* (*x*,*y*)_ and in the contrast frame *I*_*contrast* (*x*,*y*)_. Only if

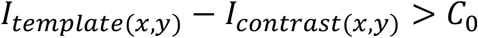

then

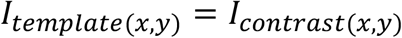

While increasing threshold *C*_0_ reduces noise, it can also lead to rejection of real movements of the fly. To optimize *C*_0_, we tested noise levels in our images by analyzing a three-hour video with dead flies. In the test, 30 pairs of consecutive frames were randomly chosen from the video and the differences between their corresponding grayscale pixel values were calculated. The distribution of the differences, stemming from noise, is shown in Figure S1C. Based on this distribution, we set *C*_0_=10, which excludes 99.99% noise-related changes of grayscale values.

#### Extracting fly shape

To extract the shape of flies in a frame, the frame is compared with the background. If a given pixel is darker on this frame than on the background frame, with the difference of grayscale being greater than threshold *C*_0_, then this pixel is temporarily assigned to the fly. That is, for pixel at location (x, y) if

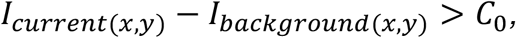

then this pixel in the current frame belongs to a fly. Despite the use of *C*_0_, some artifacts still remain in the extracted image in the form of small objects that do not belong to the fly. We eliminate these artefacts by erasing all closed objects with areas less than *C*_1_ = 20 pixels (Figure S1D), retaining only the fly silhouette (Figure S1E).

#### Feature extraction

We use normalized periphery movement (PM), core movement (CM) and centroid displacement (CD) of a fly as features for behavior classification. PM and CM are defined as the number of non-overlapping periphery and core pixels, respectively, in two consecutive frames. CD is the change of a fly’s centroid position between two frames.

#### Splitting core and periphery

To extract PM, CM and CD, we first split each fly’s body into a core and a periphery. Based on the grayscale distributions of the two parts (Figure S1F), we set the median of pixel grayscale values as the criterion to split fly body into core (darker) and periphery (lighter). This criterion makes the sizes of core and periphery to be roughly equal so that features PM and CM have equal weight in the feature space. In addition, the grayscale distribution may differ between individual animals since the light condition varies slightly across the arena. Therefore, the median value is calculated separately for each fly. In the example shown in Figure S1F, median value equals 72.

#### Centroid position

We calculate centroid position of a fly from the binary image. Suppose (*x*_1_, *y*_1_), (*x*_2_, *y*_2_), … (*x*_*n*_, *y*_*n*_) are all pixels of a fly. The centroid position is calculated from:

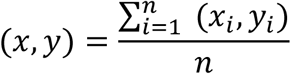

Since the tube is approximately one dimensional, when calculating centroid movement we generally ignore movements perpendicular to the long axis of the tube.

Noise may slightly change the centroid position even when a fly is stationary. Figure S1G shows the distribution of such centroid displacements caused by noise. Based on this distribution, we set 0.pixel length to be the minimum actual displacement, that is, displacements smaller than 0.5 pixel are ignored. As a result of applying this threshold, 99.66% of such false displacements are eliminated.

#### Feature normalization

. Since PM and CM both represent areas (number of pixels in area), while CD represents distance, we take the square root of PM and CM to make the dimensions of the features homogeneous. In addition, fly size varies between individuals and across experimental settings. To facilitate comparison of data in feature space, we therefore normalize PM, CM and CD of each fly with a scale parameter SP equal to the square root of the area of that fly. Thus, the final form of normalized features are

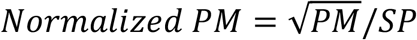

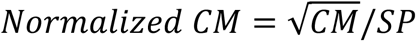

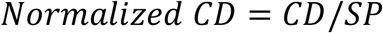

#### Orthogonality of features

In *k*NN classifier, we use Euclidean distance to measure distance in feature space between samples. Usually orthogonal features are used for this metric. By applying principal component analysis (PCA) (Jolliffe, 2002) on training data, we converted raw features (normalized PM, CM and CD) into three uncorrelated orthogonal vectors as new features. We then compared performance of the *k*NN classifier with the orthogonal features and the raw features. Based on results from 10-fold cross validation (Bishop, 2007; McLachlan, Do, & Ambroise, 2005), we found that for *k* value in *k*NN varying from 1 to 50, using additional orthogonal features does not help improve the accuracy of the classifier (Figure S1H). Since the raw features have more biophysical meaning than orthogonal features and allow us to track differences between behaviors, we opted to use normalized PM, CM and CD as features for classification.

#### Videos for Training and evaluating the kNN classifier

To construct the classifier, we visually identified 9322 frames of grooming, 9930 frames of locomotion and 5748 frames of resting from video of 20 different flies. Frames were then mapped onto the 3 dimensional PM-CM-CD feature space and used as the training set for the *k*NN classifier.

To evaluate accuracy of the classifier, we first picked a total of 15 flies from three 8 hour videos, and manually verified the accuracy of grooming events identified by our algorithm. From these videos, we randomly selected ~30 minutes video of each fly (~450 minutes in total) and manually scored all grooming events in these selected videos to identify grooming events missed by our algorithm.

#### Description of locomotion and rest behavioral classes

Since the goal of this study was a general exploration of grooming rather than a detailed classification of all fly behaviors, behaviors with body centroid movement are approximated as locomotion. For instance, feeding as measured by the amount of time spent in contact with food was classified as locomotion. Because the fly does not frequently move its body during feeding, feeding only accounts for ~1-3% in locomotion. As a result, this approximation does not significantly impact our estimation of locomotion and contributes to a considerable speed-up of analysis.

Exceptions: In Figures 4E and 5G, we explore temporal correlation between grooming and contact with food. In these figure panels only, we treated food contact separately and not as a form of locomotion. Close proximity, a body length or less, to food for >3 seconds was taken as a proxy for feeding behavior.

Except for Figures 5G and S5, rest is defined as a lack of grooming or locomotion behavior. In Figure 5G and S5, sleep is isolated from rest and described as prolonged (> 5 minutes) rest bouts. Rest other than sleep are denoted as short rest.

### Data analysis

Figure 4B: Locomotion and grooming for one day were binned every minute. Autocorrelation of each behavior is calculated at lags from 0 to 240 minutes by step of one minute. Data shown in figure is an average of 10 flies.

Figure 4C, Figure S2D: To measure periodicity in locomotion and grooming recordings, we applied the Lomb-Scargle periodogram (S. Lazopulo, Lopez, Levy, & Syed, 2015; Scargle, 1982) to time-series that were binned into 3-minute periods.

### Statistics

No sample size estimation was performed when the study was being designed. Unless otherwise specified, quantitative experiments with statistical analysis have been repeated at least three times independent. Exclusion of data applies to flies which are physically damaged (for example, broken wings or legs), physically confined (for example, trapped by condensation inside tubes), or dead during experiments. For testing statistical significance of differences between groups, we first tested the normality of data by one-sample Kolmogorov-Smirnov test. Two-sample F-test is applied for equal variances test. Samples with equal variances are compared with two-sample t-test. Satterthwaite’s approximation for the effective degrees of freedom is applied for samples with unequal variances. Results were expressed as mean ± s.d., unless otherwise specified. *p<0.05, **p<0.01, ***p<0.001 were considered statistically significant.

## Discussion

Grooming continues to be one of the least understood *Drosophila* behaviors, possibly due to the technical challenges of detecting grooming events in this small insect. Early work describing fly grooming relied on manual scoring (Connolly, 1968; Szebenyi, 1969; Tinbergen, 1965), which imposes significant limitations on the length of events that can be detected, fidelity and objectivity of detection, and the level of detail that can be extracted from the data. Despite such limitations, these initial studies made a number of noteworthy observations. Szebenyi delineated all the major modes of fly grooming and suggested that repetitive grooming actions may closely follow a preset sequence (Szebenyi, 1969). A subsequent study in the blowfly offered a more refined mechanistic picture of insect grooming by proposing that the sequential actions form a hierarchical structure (Richard & Dawkins, 1976). Combining modern computational and genetic tools, an elegant study in *Drosophila* recently confirmed these previous hypotheses (Seeds et al., 2014). That fruit flies may groom spontaneously in the absence of any apparent stimulus has also been previously suggested (Connolly, 1968; Tinbergen, 1965). Consistent with this, our work provides evidence that fruit flies groom as part of their daily repertoire of internally programmed behaviors and often without any obvious external stimulus. Our analysis revealed that, while grooming over a period of minutes appears to be spontaneous and unstructured, over a period of hours this behavior is temporally structured by the fly circadian clock, with peaks in grooming activity around dawn and dusk. The study also identifies transcription factors CLOCK and CYCLE as critical molecular components that control the amplitude of programmed *Drosophila* grooming.

Machine-learning is increasingly gaining popularity due to its applicability to virtually any problem involving pattern classification, including in studies aimed at deconstructing stereotyped behavior in the fruit fly (Branson et al., 2009; Kabra et al., 2013; Kain et al., 2013; Mendes et al., 2013; Valletta, Torney, Kings, Thornton, & Madden, 2017). Similar to these recent efforts, we constructed a computational pipeline incorporating elements of machine learning to automatically identify grooming events in video recordings of behaving flies. Our approach relies, in particular, on a supervised *k*-nearest neighbors algorithm to broadly classify behavior into grooming, locomotion and rest (Figure 2). Application of additional optional filters yields approximate data on feeding and sleep (Figure 4D, Figure 5G). While previous methods offer important details on different modes of grooming (Seeds et al., 2014), leg movements (Kain et al., 2013; Mendes et al., 2013), and fly-fly interactions (Branson et al., 2009; Kabra et al., 2013) from short videos, they demand prohibitive set-up time and computational resources for interpreting multi-day recordings. The method presented here offers less detail on modes of grooming, but can instead readily dissect circadian time-scale recordings into three-five behavioral classes on a typical personal computer.

The apparatus used in this method (Figure 1) also offers a number of advantages over current ones. First, most items used in the apparatus are standard in a typical fly circadian experiment, significantly lowering technical hurdles for other investigators to carry out similar studies. Most current grooming methods require specialized equipment for fly stimulation and detection (Seeds et al., 2014), elaborate optics, and multiple CCD cameras (Kain et al., 2013), or pre-labeled flies and a specific form of fluorescence microscopy (Mendes et al., 2013). Second, our apparatus can simultaneously monitor up to ~20 flies, while the existing approaches, though offering higher-resolution data, can monitor only one animal at a time. The scalability and high-throughput nature of our platform should appeal to investigators interested in, for example, large-scale genetic studies to identify mechanisms that differentially affect grooming, locomotion and rest (King et al., 2016). Finally, the flies in our apparatus are allowed to move freely over a distance roughly 10 times their body length and still remain in the camera’s field of view. Apparati used in other studies either constrain flies by a tether (Kain et al., 2013; Seeds et al., 2014) or permit limited visualization of behavior over short distances (Mendes et al., 2013). The relative freedom of mobility, access to food, and long time-scales of observation offered by our apparatus thus facilitate analysis of basal, internally programmed behavior.

These properties make our platform amenable to addressing questions of biological relevance, such as the importance of grooming behavior, its temporal regulation, dependence on the circadian timekeeping system, and relationship to stress. First, we found that flies consistently devote a significant fraction of time to grooming behavior during periods of locomotor activity (30%), and surprisingly, that grooming behavior is observed even during periods of reduced locomotor activity (Figure 3A). This suggests that the benefits of grooming outweigh the caloric resources expended and the resulting interruption of rest. Second, we show that daily grooming behavior, as measured by length of time spent grooming, varies less between individual flies than does locomotor activity (Figure 3E). Both of these findings underscore the hypothesis that daily grooming is a fundamental behavior of *Drosophila*.

A few recent studies (Hampel et al., 2015; Phillis et al., 1993; Seeds et al., 2014) have shown that fly grooming can be directly induced by peripheral stimuli, and there has been considerable progress toward identifying the behavioral and neural aspects of such stimulus-induced grooming. However, programmed grooming, or grooming in the absence of a macroscopic stimulus, remains relatively understudied in *Drosophila*. To our knowledge, the existence of programmed grooming, first proposed in the mid 60’s, still remains unreported.

Data from this study suggest that a significant portion of daily fly grooming is driven by internal programs. Flies in our experiments are active for ~34% of the time within a 24-hour period, during which they mostly engage in grooming, locomotion and feeding. Behavioral analysis shows that, like locomotion and feeding, grooming behavior is modulated by oscillations of the circadian clock (Figure 4). This finding raised the possibility that the observed grooming was stimulated by rhythms in contact with food or locomotor activity. However, closer examination revealed that peak in feeding activity is separated by several hours from peaks in grooming (Figure 4) and, in most cases (control and *per^0^* flies) amount of grooming remained relatively unchanged even when flies did not have access to food (Figure 5). Similarly, grooming and locomotor peaks are temporally well separated (Figure 4) and detailed examination also revealed differences in kinetic parameters underlying bout lengths and temporal patterns of grooming and locomotion (Figure 3). Additionally, genetic modifications and altered nutrient conditions resulted in contrasting changes in grooming, locomotion, and feeding (Figure 5, Figure S4). Finally, comparison of grooming in light vs. dark revealed no major differences in the fraction of daily time flies spent grooming (Figure 4D). These results together suggest that the majority of grooming events detected in our experiments are not triggered by external stimuli such as light, food, and locomotor movements. Rather, internal regulatory mechanisms, independent of external stimuli, likely drive this programmed behavior.

Multi-day recordings of wild-type flies in constant darkness showed 24-hour rhythms in daily grooming patterns. Furthermore, these rhythms were shifted appropriately in the canonical clock mutants *per^L^* and *per^S^* and abolished in the arrhythmic *per^0^* flies (Figure 4). These data support a regulatory model in which timing of programmed grooming behavior is orchestrated by the circadian clock. Notably, since these genetic perturbations did not significantly affect the amount of grooming (Figure 4D), our results suggest that the primary role of the clock is to organize the behavior in time without influencing the total time flies dedicate to grooming.

Intriguingly, two other circadian mutations, *cyc^01^* and *clk^Jrk^*, increased the proportion of daily time flies spend grooming (Figure 5A, B). *cyc^01^* flies also showed increased grooming under conditions of nutrient shortage, while *clk^Jrk^* flies showed decreased grooming under the same conditions. Importantly, neither change in grooming was observed in wild-type or *per*^0^ flies (Figure 5C), implying that the changes in grooming level are not due to circadian defects. Instead, the data imply that clock-independent but *cyc-*and *clk-*dependent pathways regulate the amount of programmed grooming behavior under normal conditions, in response to starvation, and potentially in response to other changes in the insect’s internal homeostasis.

Since both locomotion and short rest increase under starvation conditions (Figure 5G, Figure S5), it is plausible that in such situations, obtaining food is more important for survival than grooming and sleep. It may benefit the animal to have a mechanism that adjusts behavioral output to divert energy towards foraging, with *cyc* and *clk* or their products playing important roles in this regulation. This would be consistent with our observations of WT strains in starvation conditions, wherein the amount of programmed grooming remains constant despite dramatic changes in locomotion and sleep. It would also be consistent with our observations of *cyc^01^* and *clk^Jrk^* flies, which show altered grooming when nutrients are unavailable, presumably due to defective regulation of behavioral output. Differences in starvation-induced changes between *cyc^01^* and *clk^Jrk^* flies suggest an additional mechanistic detail regarding the *cyc-*and *clk-*mediated pathways. When subjected to sleep deprivation, *cyc^01^* but not *clk^Jrk^* flies, dramatically lower expression of heat-shock genes, and show excessive homeostatic rebound (Shaw et al., 2002). In the present context, these prior data raise the possibility that heat-shock genes might also be part of the *cyc^01^* and *clk^Jrk^* dependent grooming response pathways that are activated by starvation.

Finally, why are flies innately programmed to groom? The present study does not directly address this important question, but given that microscopic pathogens can sporulate on the fly cuticle and eventually infect the insect (Leger, Wang, & Fang, 2011), persistent grooming may serve as a first line of defense against such attack. Thus, the immune system may constitute another internal program, similar to the *cyc* and *clk*-controlled mechanisms, that drives fly grooming; if so, we hypothesized that mutants with defective immune response may exhibit altered grooming behavior (Lemaitre et al., 1995; Michel, Reichhart, Hoffmann, & Royet, 2001). Consistent with this, we found that grooming was reduced in the immune deficient *imd* mutant (Figure S6A), though a second immune deficient strain lacking a member of the Toll pathway (*PGRP-SA^seml^)* showed only a modest decrease (Figure S6A). Further studies are required to clarify these initial results and elucidate the biological function of programmed grooming in *Drosophila*.

Together, the data provide strong supporting evidence for programmed grooming in *Drosophila* and suggest that this innate behavior is driven by two distinct sets of regulatory systems. The circadian system temporally segregates undulations in grooming from those of other essential behavioral outputs like feeding and sleep. Circadian coordination of grooming underscores a previously under-appreciated importance of this behavior in the daily routine of the fruit fly. The second regulatory system adjusts the level of grooming relative to other behaviors. This set of regulation likely confers adaptability on the animal by allowing it to up-or downregulate grooming as necessitated by internal and external conditions. The dual control mechanism of grooming proposed here is highly reminiscent of the two-process framework–-circadian and homeostatic–-that is widely used in understanding sleep regulation (Borbély, 1982). Although this work has not demonstrated grooming is under homeostatic control, future studies could be aimed at better characterizing the nature of the non-circadian regulatory system of fly grooming.

In summary, we present here a new platform to detect innate grooming behavior simultaneously and for days at a time in multiple individual fruit flies. The apparatus can be assembled easily, and the accompanying analytics is available publicly. Utilizing this platform, we report several mechanisms that are potentially responsible for driving the timing and level of programmed grooming in *Drosophila*. We also suggest future experiments that through use of this platform can lead to deeper understanding of the underlying biology of grooming and its relation to other essential fly behaviors.

## Acknowledgements

This work was partially supported by the National Science Foundation under grant IOS-1656603 to S.S. and by National Institutes of Health grants R01GM105775 and R01AG045842 to M.M.S.H. The authors are grateful to Michael Young and William Ja for providing fly strains, Juan Lopez and Manuel Collazo for technical support and Stanislav Lazopulo and Andrey Lazopulo for suggestions and assistance with experiments. We thank Alan Li and Gadi Trocki for helpful comments on the manuscript.

---

**Supplementary Figure S1:**
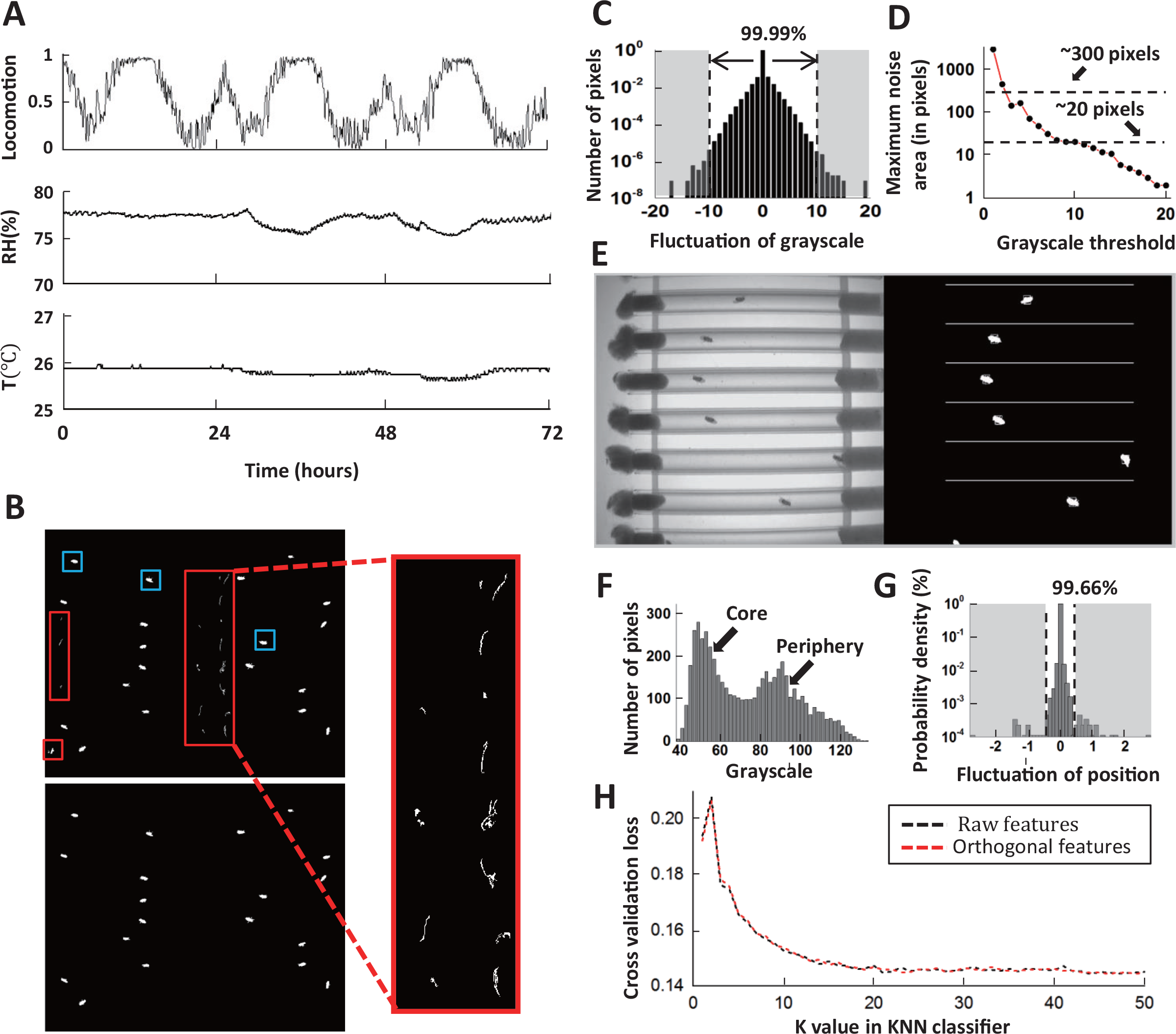
Grooming tracking algorithm. (A) Locomotion (fraction of time spent), relative humidity (RH), and temperature (T) for 3 days in constant darkness (DD) conditions. Data are binned in five minutes. (B) Binary images after background subtraction. If the background frame is not updated frequently (typically every 1000 seconds), both food debris (red boxes) and flies (blue boxes) may be identified as moving objects in a background-subtracted image (top, left and expanded view). The problem is rectified (bottom, left) when the background frame used is closer in time (<1000 seconds apart) to the image of interest. (C) The distribution of grayscale fluctuations in the absence of mobile flies. A cutoff of grayscale value change *C*_0_ = 10 rules out > 99.99% of fluctuations. (D)Maximum area (pixels) of a closed object generated by noise when different thresholds *C*_0_ are applied. A choice of *C*_0_ = 10 rejects objects larger than 20 pixels without affecting identification of flies which have a typical area of ~300 pixels in our studies. (E) An example 8-bit frame (on left) and its corresponding background-subtracted binary image showing identified flies. (F) Grayscale value distribution of pixels belonging to 20 individual flies. Two regions are clearly seen: the left region with peak around 50 represents the core of the flies and the right region with peak around 90 represents their periphery. (G) Variations in the center position of a stationary fly. The minimum displacement that represents a true fly center movement is 0.5 pixel length in our experiment, a requirement that excludes 99.66% of false displacements. (H) The cross validation loss of *k*NN classifier at different *k* values. No significant difference between using raw features (black) or PCA-derived orthogonal features (red). Loss decreases with increasing *k* values, slowing down for *k*≈10. The loss function shown here is the averaged error of 10-fold cross validation in behavioral classification. The validation was performed on 25000 frames from video of 20 flies.

**Supplementary Figure S2:**
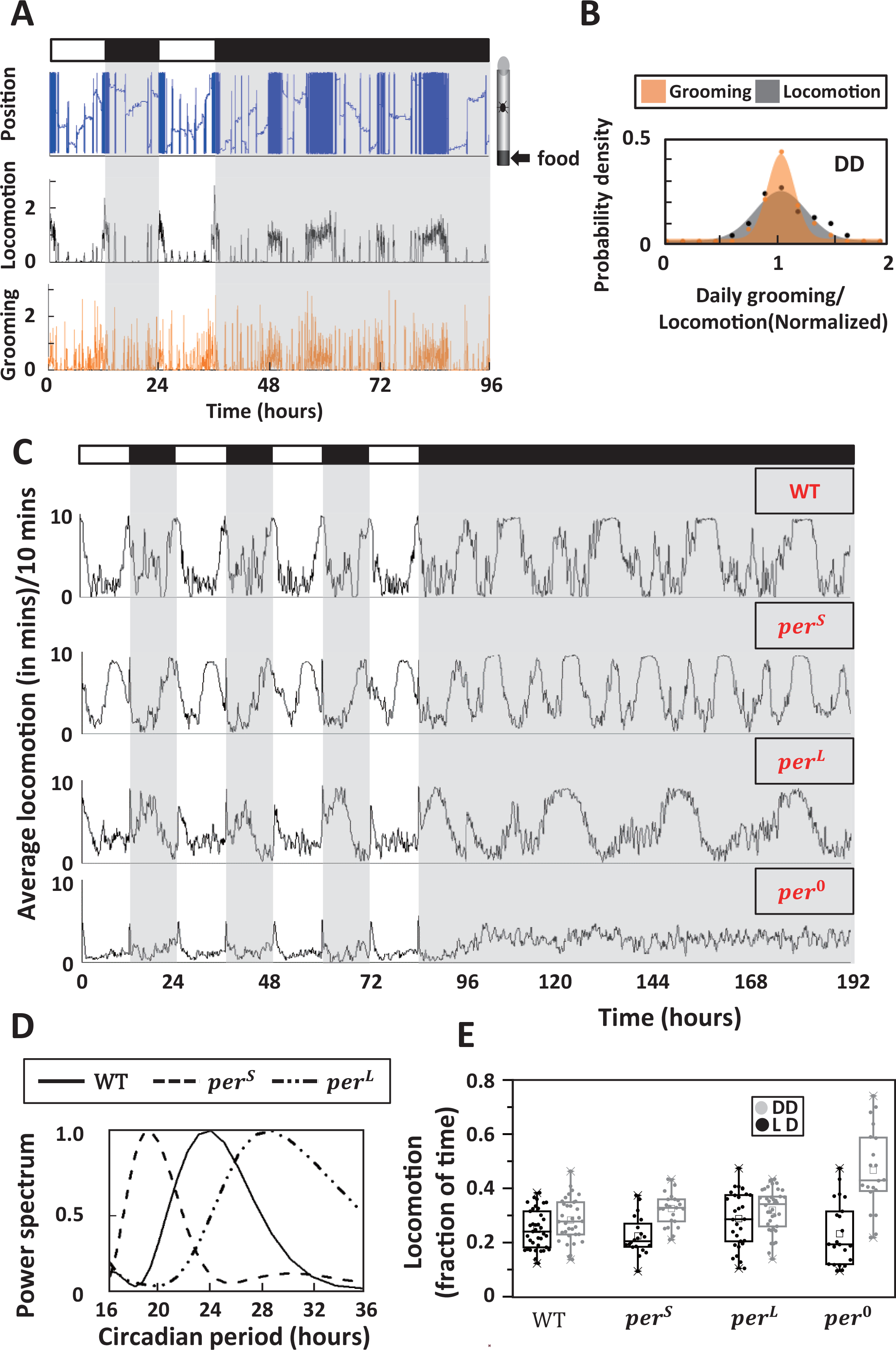
Circadian regulation on locomotion. (A) Position (top row), locomotion level (middle) and grooming level (bottom) of a single fly during two days in LD followed by two days in DD conditions. Locomotion and grooming are shown in terms of the amount of time (in minutes) spent by the fly in the two activities. The data are plotted in 5 min bins. White/black bars indicate light/dark conditions, respectively. (B) Inter-individual differences of daily grooming and locomotion in constant darkness. Distributions of normalized individual grooming and locomotion (amount of daily grooming/locomotion of individuals divided by population average) are fitted to normal distribution. Variation in daily grooming time among individuals is significantly less than the variation in locomotion with the standard deviation of grooming being 0.16 and that of locomotion being 0.25. N=34 wild-type flies. (C) Locomotor activity (in 10 minute bins) of WT and clock mutants during four days in LD cycle followed by four days in DD cycles. Both activities are population averages. N=8 WT, 8 *per*^*S*^, 10 *per*^*L*^, 10 *per*^0^. (D) Average power spectra of wild-type, *per*^*S*^ and *per*^*L*^ locomotion in DD. N=34 wildtype, 23 *per*^*S*^, 38 *per*^*L*^. (E) Daily time spent in locomotion by WT and clock mutants under LD and DD cycles. In most cases, locomotion time increases under constant darkness.

**Supplementary Figure S3:**
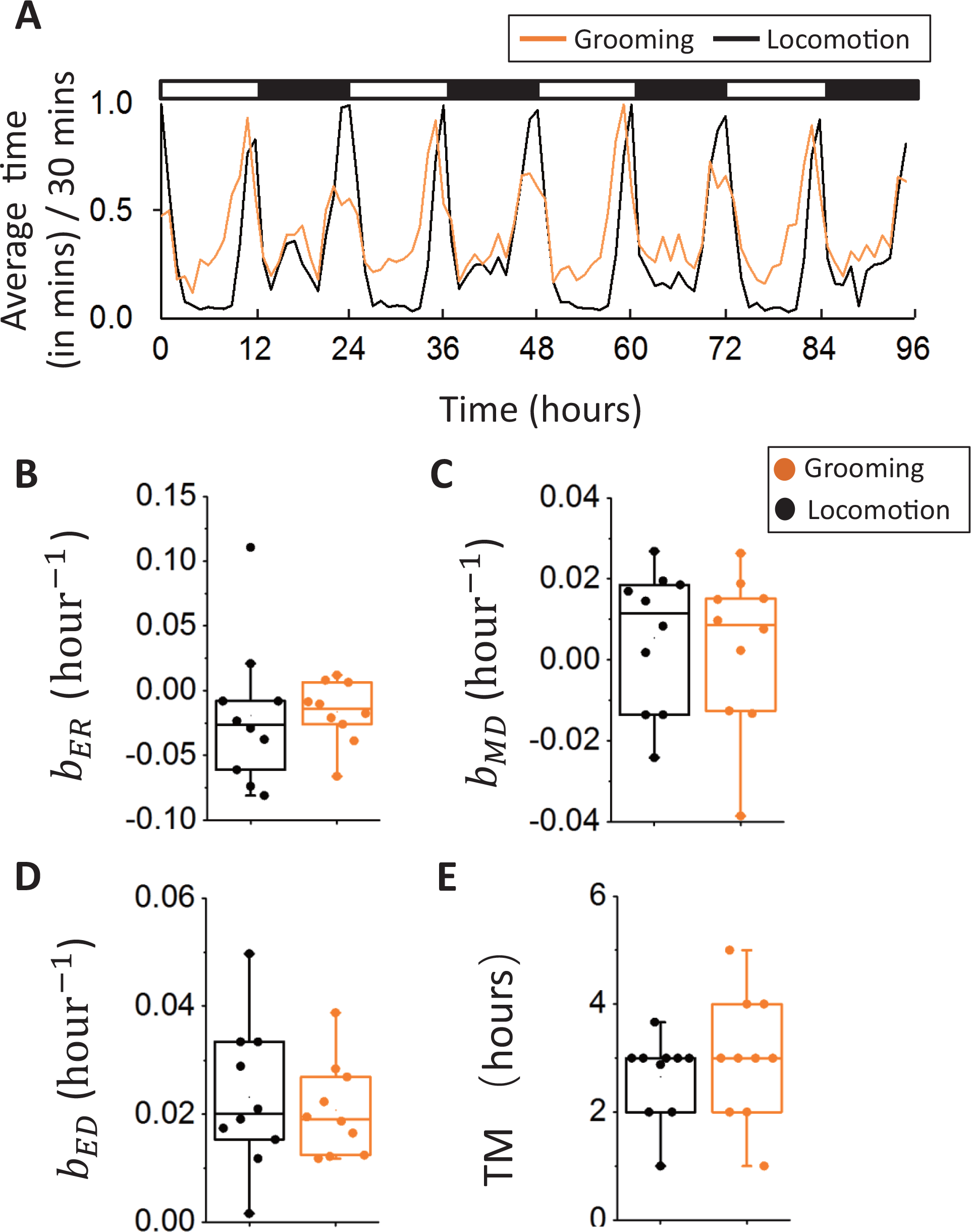
(A) Normalized average amount time spent in grooming (orange) and locomotion (black) during four days in LD. Each behavior time series is normalized by its maximum. Change of grooming between day and night, especially the change from night to morning, is smaller than the corresponding change in locomotion. This difference between grooming and locomotion indicates a small increasing rate parameter (*b*_*MR*_) for grooming. N=10 WT flies. (B)(C) (D) (E) Rate parameters *b*_*MD*_, *b*_*ER*_, *b*_*ED*_ and duration of morning peaks (TM) do not show significant differences between grooming and locomotion.

**Supplementary Figure S4.**
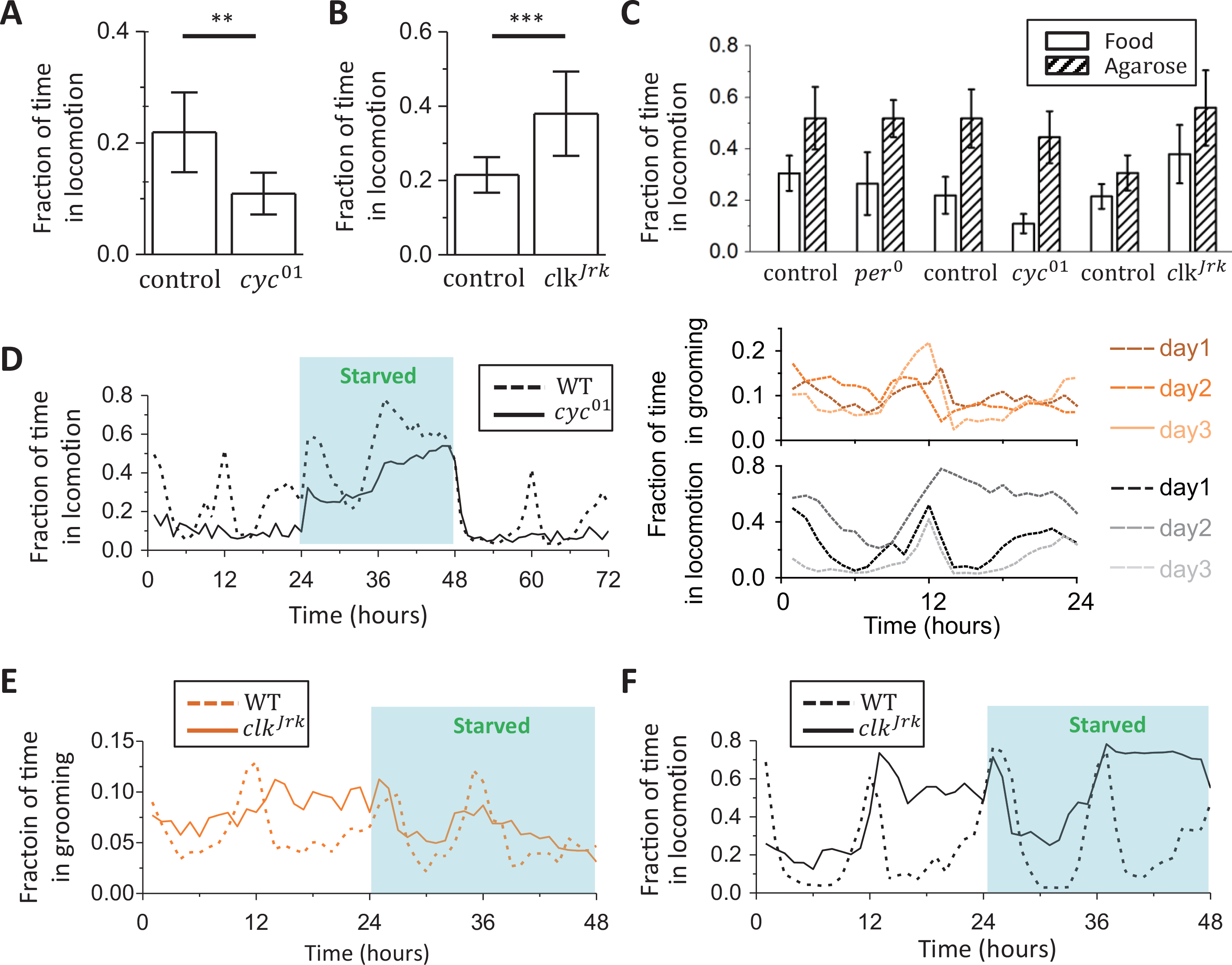
(A), (B) Fraction of time spent daily in locomotion by *cyc^01^*, *clk^Jrk^* and their controls. *cyc^01^* flies spend less time in locomotion than control flies (p=0.0014). In contrast, *clk^Jrk^* flies dedicate more time to locomotor activity than their controls (p<0.001). (C) Fraction of time spent in locomotion in response to stress through starvation. All strains of flies show increased amount of locomotion when starved. N=18 *per*^0^ flies and 20 of control. N=18 *cyc*^01^ flies and 14 of control. N=28 *clk^Jrk^* flies and 28 of control. (D) Temporal patterns in WT (N= 10) and *cyc*^01^ (N= 10) locomotion during a 3-day 12:12 LD experiment in which flies are starved on a 1% agarose diet during the second day (shaded). Population average data plotted in one-hour bins. Flies show elevated locomotion when starved. In two panels on right, WT grooming and locomotion from individual days are plotted separately for comparison. (E), (F) Temporal patterns of (E) grooming and (F) locomotion of control (N= 10) and *clk^Jrk^* (N= 10) flies during the first 2 days of a starvation experiment. In the experiment, flies are given regular corn meal on the first day, and 1% agarose on day 2 (shaded). The data are shown in one-hour bins.

**Supplementary Figure S5.**
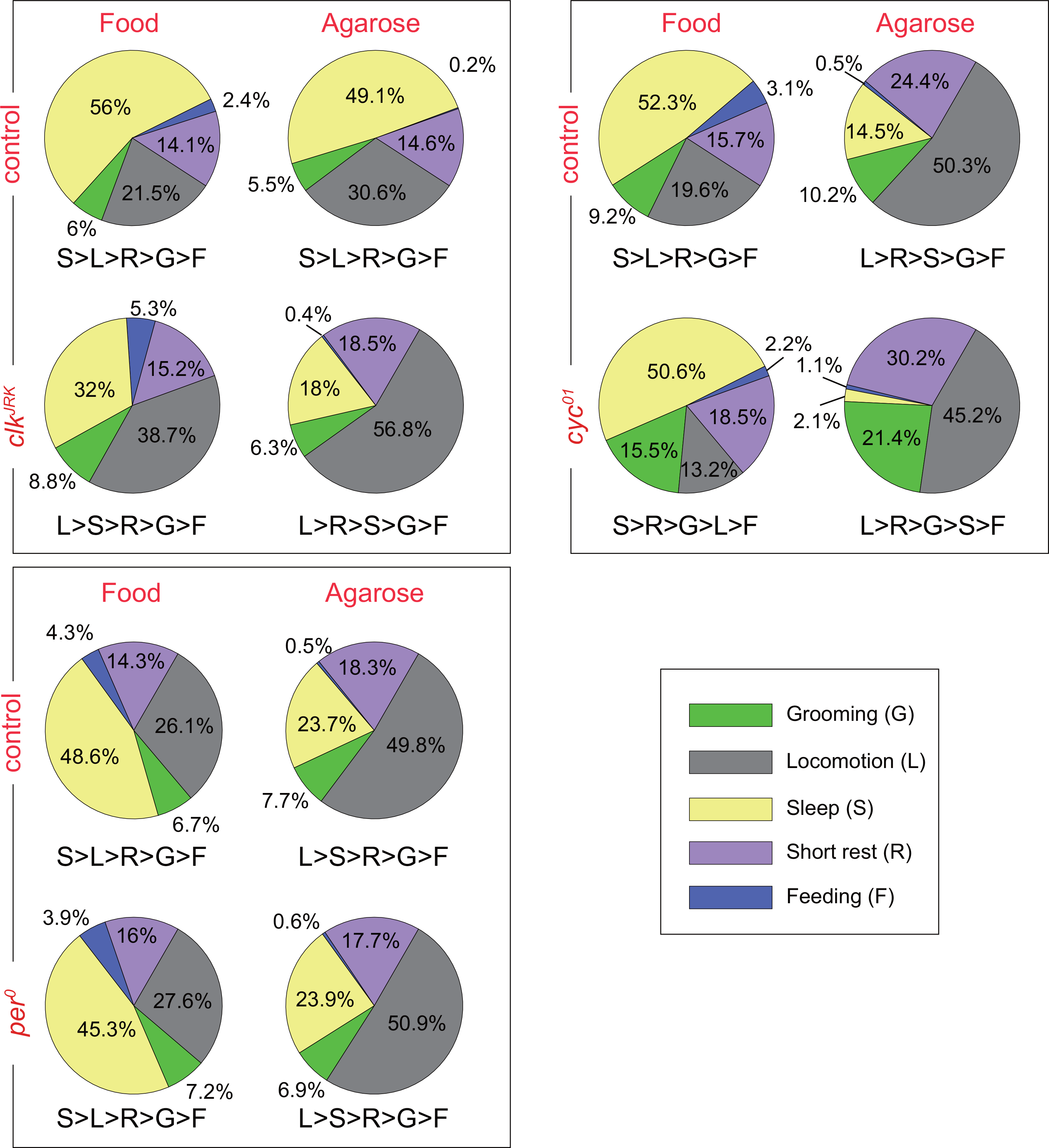
Average fraction of time flies spend in grooming (green), locomotion (gray), sleep (yellow), short rest (purple) and feeding (blue). N=17 *per^0^* flies and 20 of control. N=18 *cyc^01^* flies and 1 of control. N=25 *clk^Jrk^* flies and 28 of control. The ranked amount of time in behaviors is shown below each pie-chart, with G, L, S, R, F representing grooming, locomotion, sleep, short rest and feeding, respectively.

**Supplementary Figure S6.**
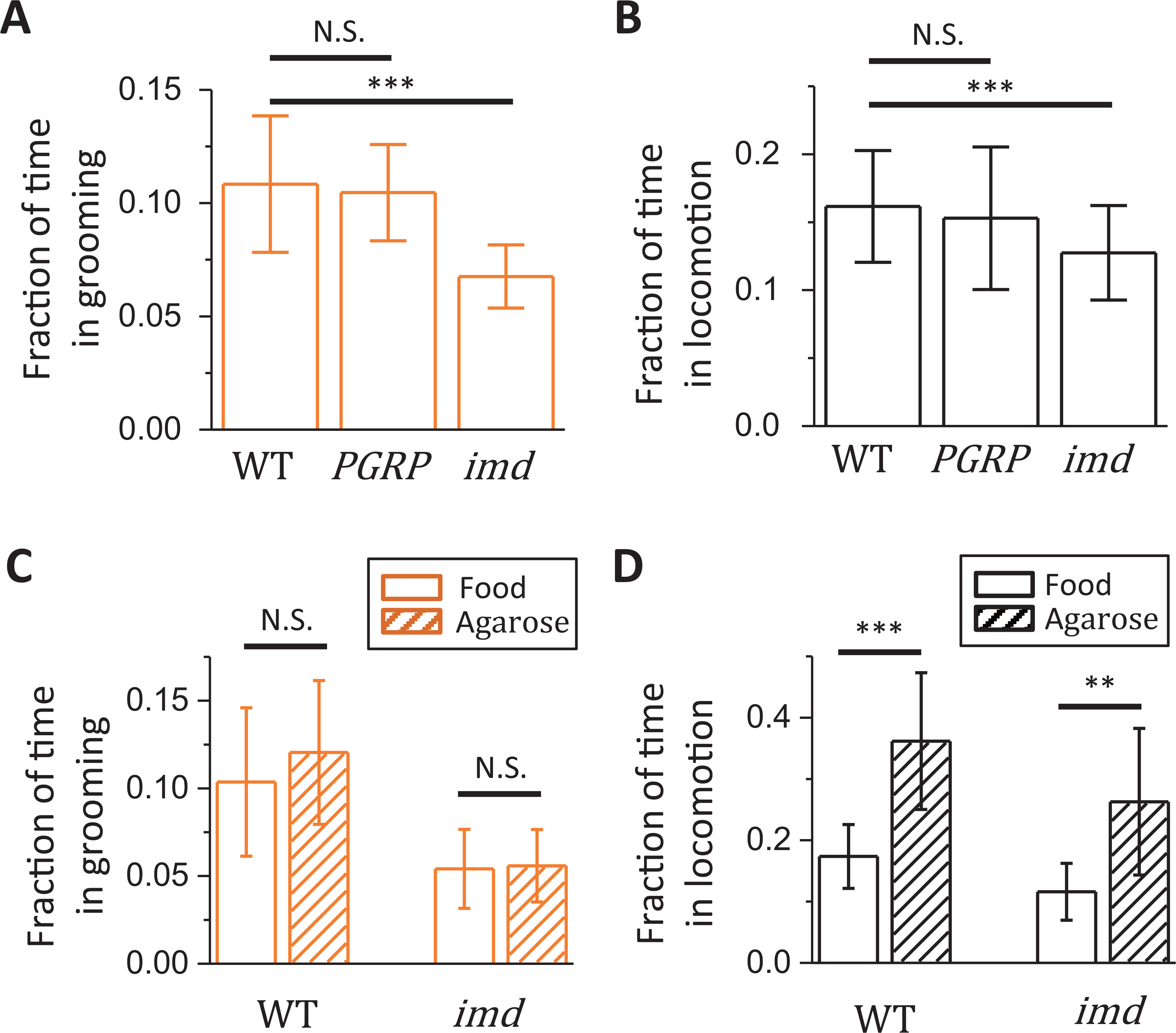
Immune systems may regulate the amount of grooming. Pathogens can infect fly through breaching the cuticle. Since one of the main function of grooming is to keep body surface clean, it is possible that grooming might work as part of immune systems. We test two mutant fly strains *imd* and *PGRP-SA*^seml^, both of which have defective immune systems. Mutants *PGRP-SA*^seml^ and *imd* are on *Oregon R* background. (A), (B) Grooming and locomotion in *imd* flies are significantly less than control flies (p<0.001 for both grooming and locomotion), while *PGRP-SA*^seml^ does not significantly affect the time spent in grooming or locomotion. This suggests that *Drosophila* grooming relies on a working immune system. The decrease in *imd* flies further suggests that this impact may be independent of the Toll pathway. (C), (D) In both *imd* and control flies, locomotion increases significantly when starved (p<0.001 for WT and p<0.01 for *imd*), without a robust change in grooming in either strain.

## Rich Media Files

**Video 1: Sample raw experimental video**

**Video 2: Sample video of grooming on head and front legs**

**Video 3: Sample video of grooming on wings and hind legs**

## References

Allada, R., & Chung, B. Y. (2010). Circadian organization of behavior and physiology in Drosophila. Annual Review of Physiology, 72, 605–24. http://doi.org/10.1146/annurev-physiol-021909-135815

Allada, R., White, N. E., So, W. V., Hall, J. C., & Rosbash, M. (1998). A mutant Drosophila homolog of mammalian Clock disrupts circadian rhythms and transcription of period and timeless. Cell, 93(5), 791–804. http://doi.org/10.1016/S0092-8674(00)81440-3

Bergman, C. M., Schaefer, J. A., & Luttich, S. N. (2000). Caribou movement as a correlated random walk. Oecologia, 123(3), 364–374. http://doi.org/10.1007/s004420051023

Bishop, C. M. (2007). Pattern Recognition and Machine Learning. Springer.

Borbély, A. A. (1982). A two process model of sleep regulation. Human Neurobiology, 1(3), 195-204.

Branson, K., Robie, A. A., Bender, J., Perona, P., & Dickinson, M. H. (2009). High-throughput ethomics in large groups of Drosophila. Nature Methods, 6(6), 451–457. http://doi.org/10.1038/nmeth.1328

Chatterjee, A., Tanoue, S., Houl, J. H., & Hardin, P. E. (2010). Regulation of Gustatory Physiology and Appetitive Behavior by the Drosophila Circadian Clock. Current Biology, 20(4), 300–309. http://doi.org/10.1016/j.cub.2009.12.055

Chen, S., Tvrdik, P., Peden, E., Cho, S., Wu, S., Spangrude, G., & Capecchi, M. R. (2010). Hematopoietic Origin of Pathological Grooming in Hoxb8 Mutant Mice. Cell, 141(5), 775–785. http://doi.org/10.1016/j.cell.2010.03.055

Connolly, K. (1968). The social facilitation of preening behaviour in Drosophila melanogaster. Animal Behaviour, 16(2), 385–391. http://doi.org/10.1016/0003-3472(68)90023-7

Crane, B. R., & Young, M. W. (2014). Interactive features of proteins composing eukaryotic circadian clocks. Annual Review of Biochemistry, 83, 191–219. http://doi.org/10.1146/annurev-biochem-060713-035644

Ferkin, M. H., Leonard, S. T., Heath, L. A., & Paz-y-Miño, C. G. (2001). Self-grooming as a tactic used by prairie voles Microtus ochrogaster to enhance sexual communication. Ethology, 107(10), 939–949. http://doi.org/10.1046/j.1439-0310.2001.00725.x

Geist, Valerius. Walther, F. (1974). The behaviour of ungulates and its relation to management. I.U.C.N. Publication, 24, 1–194.

Gilestro, G. F. (2012). Video tracking and analysis of sleep in Drosophila melanogaster. Nature Protocols, 7(5), 995–1007. http://doi.org/10.1038/nprot.2012.041

Hampel, S., Franconville, R., Simpson, J. H., & Seeds, A. M. (2015). A neural command circuit for grooming movement control. eLife, 4(September), 1–26. http://doi.org/10.7554/eLife.08758

Hart, B. L. (1988). Biological basis of the behavior of sick animals. Neuroscience & Biobehavioral Reviews, 12(2), 123–137. http://doi.org/10.1016/S0149-7634(88)80004-6

Hart, B. L., Hart, L. A., Mooring, M. S., & Olubayo, R. (1992). Biological basis of grooming behaviour in antelope: the body-size, vigilance and habitat principles. Animal Behaviour, 44(4), 615–631. http://doi.org/10.1016/S0003-3472(05)80290-8

Hawlena, H., Bashary, D., Abramsky, Z., Khokhlova, I. S., & Krasnov, B. R. (2008). Programmed versus stimulus-driven antiparasitic grooming in a desert rodent. Behavioral Ecology, 19(5), 929–935. http://doi.org/10.1093/beheco/arn046

Hendricks, J. C., Lu, S., Kume, K., Yin, J. C., Yang, Z., & Sehgal, A. (2003). Gender dimorphism in the role of cycle (BMAL1) in rest, rest regulation, and longevity in Drosophila melanogaster. Journal of Biological Rhythms, 18(1), 12–25. http://doi.org/10.1177/0748730402239673

Jolliffe, I. (2002). Principal component analysis. Wiley Online Library.

Kabra, M., Robie, A. A., Rivera-Alba, M., Branson, S., & Branson, K. (2013). JAABA: interactive machine learning for automatic annotation of animal behavior. Nature Methods, 10(1), 64–67. http://doi.org/10.1038/nmeth.2281

Kain, J., Stokes, C., Gaudry, Q., Song, X., Foley, J., Wilson, R., & de Bivort, B. (2013). Leg-tracking and automated behavioural classification in Drosophila. Nature Communications, 4, 1910. http://doi.org/10.1038/ncomms2908

Keene, A. C., Duboué, E. R., McDonald, D. M., Dus, M., Suh, G. S., Waddell, S., & Blau, J. (2010, July). Clock and cycle limit starvation-induced sleep loss in Drosophila. Current Biology. http://doi.org/10.1016/j.cub.2010.05.029

King, L. B., Koch, M., Murphy, K., Velazquez, Y., William, W. J., & Tomchik, S. M. (2016). Neurofibromin Loss of Function Drives Excessive Grooming in Drosophila. Genes| Genomes| Genetics, 6, 1083–1093. http://doi.org/10.1534/g3.115.026484

Lazopulo, A., & Syed, S. (2016). A mathematical model provides mechanistic links to temporal patterns in Drosophila daily activity. BMC Neuroscience, 17(1), 14. http://doi.org/10.1186/s12868-016-0248-9

Lazopulo, S., Lopez, J. A., Levy, P., & Syed, S. (2015). A stochastic burst follows the periodic morning peak in individual Drosophila locomotion. PLoS ONE, 10(11), 1–17. http://doi.org/10.1371/journal.pone.0140481

Leger, R. J., Wang, C., & Fang, W. (2011). New perspectives on insect pathogens. Fungal Biology Reviews, 25(2), 84–88. http://doi.org/10.1016/j.fbr.2011.04.005

Lemaitre, B., Kromer-Metzger, E., Michaut, L., Nicolas, E., Meister, M., Georgel, P., … Hoffmann, J. A. (1995). A recessive mutation, immune deficiency (imd), defines two distinct control pathways in the Drosophila host defense. Proceedings of the National Academy of Sciences, 92(21), 9465–9469. http://doi.org/10.1073/pnas.92.21.9465

McKenna, J. J. (1978). Biosocial functions of grooming behavior among the common Indian langur monkey (Presbytis entellus). American Journal of Physical Anthropology, 48(4), 503–509. http://doi.org/10.1002/ajpa.1330480409

McLachlan, G., Do, K. A., & Ambroise, C. (2005). Analyzing microarray gene expression data (Vol. 422). John Wiley & Sons.

Mendes, C. S., Bartos, I., Akay, T., Márka, S., & Mann, R. S. (2013). Quantification of gait parameters in freely walking wild type and sensory deprived Drosophila melanogaster. eLife, 2, e00231. http://doi.org/10.7554/eLife.00231

Michel, T., Reichhart, J. M., Hoffmann, J. A., & Royet, J. (2001). Drosophila Toll is activated by Gram-positive bacteria through a circulating peptidoglycan recognition protein. Nature, 414(6865), 756–759. http://doi.org/10.1038/414756a

Mooring, M. S., Blumstein, D. T., & Stoner, C. J. (2004). The evolution of parasite-defence grooming in ungulates. Biological Journal of the Linnean Society, 81(1), 17–37. http://doi.org/10.1111/j.1095-8312.2004.00273.x

Mooring, M. S., & Samuel, W. M. (1998). The biological basis of grooming in moose: programmed versus stimulus-driven grooming. Animal Behaviour, 56(6), 1561–1570. http://doi.org/10.1006/anbe.1998.0915

Owald, D., Lin, S., & Waddell, S. (2015). Light, heat, action: neural control of fruit fly behaviour. Philosopical Transactions of the Royal Society B, Biological Sciences, 370(1677), 20140211. http://doi.org/10.1098/rstb.2014.0211

Patenaude, F., & Bovet, J. (1984). Self-grooming and social grooming in the North American beaver, Castor canadensis. Canadian Journal of Zoology, 62(9), 1872–1878. http://doi.org/10.1139/z84-273

Pfeiffenberger, C., Lear, B. C., Keegan, K. P., & Allada, R. (2010). Locomotor Activity Level Monitoring Using the Drosophila Activity Monitoring (DAM) System. Cold Spring Harbor Protocols, 2010(11), pdb.prot5518. http://doi.org/10.1101/pdb.prot5518

Phillis, R. W., Bramlage, A. T., Wotus, C., Whittaker, A., Gramates, L. S., Seppala, D., … Murphey, R. K. (1993). Isolation of Mutations Affecting Neural Circuitry Required for Grooming Behavior in Drosophila melanogaster. Genetics, 133, 581–592.

Richard, & Dawkins, M. (1976). Hierachical organization and postural facilitation: Rules for grooming in flies. Animal Behaviour, 24(4), 739–755. http://doi.org/10.1016/S0003-3472(76)80003-6

Rutila, J. E., Suri, V., Le, M., So, W. V., Rosbash, M., & Hall, J. C. (1998). CYCLE is a second bHLH-PAS clock protein essential for circadian rhythmicity and transcription of Drosophila period and timeless. Cell, 93(5), 805–814. http://doi.org/10.1016/S0092-8674(00)81441-5

Sachs, B. D. (1988). The development of grooming and its expression in adult animals. Annals of the New York Academy of Sciences, 525, 1–17. http://doi.org/10.1111/j.1749-6632.1988.tb38591.x

Scargle, J. D. (1982). Studies in astronomical time series analysis. II - Statistical aspects of spectral analysis of unevenly spaced data. The Astrophysical Journal, 263, 835–853. http://doi.org/10.1086/160554

Schino, G. (2001). Grooming, competition and social rank among female primates: a meta-analysis. Animal Behaviour, 62, 265–271. http://doi.org/10.1006/anbe.2001.1750

Schino, G., Scucchi, S., Maestripieri, D., & Turillazzi, P. G. (1988). Allogrooming as a tension-reduction mechanism: A behavioral approach. American Journal of Primatology, 16(1), 43–50. http://doi.org/10.1002/ajp.1350160106

Schlichting, M., Menegazzi, P., Lelito, K. R., Yao, Z., Buhl, E., Dalla Benetta, E., … Shafer, O. T. (2016). A Neural Network Underlying Circadian Entrainment and Photoperiodic Adjustment of Sleep and Activity in Drosophila. The Journal of Neuroscience, 36(35), 9084–96. http://doi.org/10.1523/JNEUROSCI.0992-16.2016

Seeds, A. M., Ravbar, P., Chung, P., Hampel, S., Midgley, F. M., Mensh, B. D., & Simpson, J. H. (2014). A suppression hierarchy among competing motor programs drives sequential grooming in Drosophila. eLife, 3, e02951. http://doi.org/10.7554/eLife.02951

Seyfarth, R. M. (1977). A model of social grooming among female monkeys. Journal of Theoretical Biology, 65(4), 671–698. http://doi.org/10.1016/0022-5193(77)90015-7

Shaw, P. J., Tononi, G., Greenspan, R. J., & Robinson, D. F. (2002). Stress response genes protect against lethal effects of sleep deprivation in Drosophila. Nature, 417(6886), 287–291. http://doi.org/10.1038/417287a

Sproull, R. F. (1991). Refinements to nearest-neighbor searching in k-dimensional trees. Algorithmica, 6(1), 579–589. http://doi.org/10.1007/BF01759061

Spruijt, B. M., van Hooff, J. A., & Gispen, W. H. (1992). Ethology and neurobiology of grooming behavior. Physiological Reviews, 72(3), 825–852.

Stoleru, D., Peng, Y., Agosto, J., & Rosbash, M. (2004). Coupled oscillators control morning and evening locomotor behaviour of Drosophila. Nature, 431(7010), 862–868. http://doi.org/10.1038/nature02926

Szebenyi, A. L. (1969). Cleaning Behaviour In Drosophila Melanogaster. Animal Behaviour, 17(1), 641–651. http://doi.org/10.1016/S0003-3472(69)80006-0

Thiessen, D. D., Graham, M., Perkins, J., & Marcks, S. (1977). Temperature regulation and social grooming in the Mongolian gerbil (Meriones unguiculatus). Behavioral Biology, 19(3), 279–88. http://doi.org/10.1016/S0091-6773(77)91579-6

Tinbergen, N. (1965). Animal behavior. Time Incorporated.

Valletta, J. J., Torney, C., Kings, M., Thornton, A., & Madden, J. (2017). Applications of machine learning in animal behaviour studies. Animal Behaviour, 124, 203–220. http://doi.org/10.1016/j.anbehav.2016.12.005

Walther, F. R. (1984). Communication and expression in hoofed mammals. Indiana University Press.

Xu, K., Zheng, X., & Sehgal, A. (2008). Regulation of feeding and metabolism by neuronal and peripheral clocks in Drosophila. Cell Metabolism, 8(4), 289–300. http://doi.org/10.1016/j.cmet.2008.09.006

Yanagawa, A., Guigue, A. M. a, & Marion-Poll, F. (2014). Hygienic grooming is induced by contact chemicals in Drosophila melanogaster. Frontiers in Behavioral Neuroscience, 8, 254. http://doi.org/10.3389/fnbeh.2014.00254

